# Maternal effects and environmental filtering shape seed fungal communities in oak trees

**DOI:** 10.1101/691121

**Authors:** Tania Fort, Charlie Pauvert, Amy E. Zanne, Otso Ovaskainen, Thomas Caignard, Matthieu Barret, Stéphane Compant, Arndt Hampe, Sylvain Delzon, Corinne Vacher

**Affiliations:** BIOGECO, INRA, Univ. Bordeaux, 33615 Pessac, France; George Washington University, Biological Sciences Department, 800 22^nd^ St., Washington DC 20052, USA; Organismal and Evolutionary Biology Research Programme, P.O. Box 65, 00014 University of Helsinki, Finland; Center for Biodiversity Dynamics, Department of Biology, Norwegian University of Science and Technology, 7491 Trondheim, Norway; IRHS, Agrocampus-Ouest, INRA, Université d’Angers, SFR4207 QuaSaV, 49071 Beaucouzé, France; AIT Austrian Institute of Technology GmbH, Center for Health & Bioresources, Bioresources Unit, Konrad-Lorenz Straße 24, 3430 Tulln, Austria; Univ. Bordeaux, INRA, BIOGECO, Pessac France

**Keywords:** Microbial ecology, disease ecology, community genetics, ecological networks, joint species distribution models, endophyte, host-parasite interaction, vertical transmission

## Abstract

- Trees, as foundation species, play a pivotal role in the species interaction networks that constitute forest ecosystems. From the seed stage, they interact with microbial communities that affect their growth, health and fitness. Despite their eco-evolutionary importance, the processes shaping seed microbial communities in natural forests have received little attention.
- To unravel these processes, we analyzed the microbial communities of seeds collected in populations of sessile oak (*Quercus petraea*) growing along elevation gradients. We focused on the fungal communities as this group includes seed pathogens. Ecological processes shaping the communities were quantified using joint species distribution models.
- Fungi were present in all seed tissues, including the embryo. Fungal communities differed significantly among oak populations along the elevation gradients, and among mother trees within the same population. These maternal effects remained significant after seed fall, despite colonization by fungal species on the ground. Associations between tree pathogens and their antagonists were detected in the seeds.
- Our results demonstrate that both maternal effects and environmental filtering shape seed microbial communities of sessile oak. They provide a starting point for future research aimed at identifying the seed extended phenotypic traits that influence seed dispersal and germination, and seedling survival and growth across environments.

## Introduction

Seeds are colonized by a wide range of microorganisms that positively and negatively influence plant growth, health and fitness. They associate with many endophytic and epiphytic microorganisms that foster the growth of seedlings and protect them against natural enemies (Baker & Smith, 1966; Nelson, 2004; Links, *et al.* 2014). For example, the seed-borne endophytic fungus *Epichloë festucae* limits pest attacks and pathogen development by producing alkaloids (Clay & Schardl, 2002; Faeth, 2002). Seeds are also colonized by pathogens that reduce germination rate (Nelson, 2017) and seedling survival (Kremer, 1987; Gilbert, 2002). For example, the fungal pathogen *Ciboria batschiana*, the causal agent of acorn black rot, can damage up to 80% of acorns when conditions are wet (Prochazkova *et al.*, 2005). By affecting the early stages of the plant life cycle, seed pathogens have significant impacts on natural ecosystems, including forests in which they shape tree species diversity and spatial structure (Janzen, 1970; Gilbert, 2002). Understanding how pathogens and other microorganisms are acquired and transmitted to the seeds of foundation tree species (i.e. tree species that structure and stabilize forest communities and ecosystem processes; Whitham *et al.*, 2006), and how these microorganisms associate to regulate seed germination and seedling development, is thus crucial to our ability to predict and manage the regeneration of forest ecosystems.

Up to now, processes of seed microbiota assembly have received more attention in crop plants than in natural ecosystems. In crops, microorganisms are transmitted by the mother plant to its seeds at the floral and during early seed development stages, through vascular tissues or contact between vegetative and reproductive organs (Maude, 1996). They can also be transmitted from the pollen of the father plant, insect vectors, or bioaerosols (Escobar Rodríguez *et al.*, 2018; Frank *et al.*, 2017). Once seeds fall on the ground, epiphytic microbial communities coalesce with microbial communities of litter and upper soil (Rillig *et al.*, 2015). Germinating seeds then release molecules that attract soil microbes, surrounding themselves with a microbiologically active soil area called the spermosphere (Nelson, 2004; Schiltz *et al.*, 2015). The emergence of the plant radicle creates cracks in the seed tegument, enabling microbes to colonize internal tissues (Nelson *et al.*, 2018). These events lead to intense biotic interactions among microorganisms (Nelson, 2004) and drastic changes in seed microbiota composition and function (Ofek *et al.*, 2011; Yang *et al.*, 2017; Torres-Cortés *et al.*, 2018). Recent studies suggest that seed colonization by soil microorganisms represents the most influential microbial acquisition for seedling growth and health (Nelson *et al.*, 2018).

The seed microbiota is therefore a dynamic entity that is shaped, like all ecological communities, by four fundamental processes: selection by the abiotic environment (environmental filtering) and biotic interactions (biotic filtering), dispersal, ecological drift and evolutionary diversification (Vellend, 2010; Nemergut *et al.*, 2013; Ovaskainen *et al.*, 2017; Zhou & Ning, 2017). Unraveling these assembly processes is particularly important because they govern seed and seedling extended phenotypes (defined as the diversity and composition of associated communities; Whitham *et al.*, 2006), as well as the performance and eventually fitness of the plant host (Compant *et al.*, 2010; Truyens *et al.*, 2015; Brader *et al.*, 2017). Some processes are deterministic and depend on taxon-specific functional traits (Minard *et al.*, 2019), such as the response to selection. Other processes are partly or purely stochastic (Zhou & Ning, 2017; Rezki *et al.*, 2018) and generate divergences among communities occupying identical environments (Chase & Myers, 2011). The dispersal process is particular in the case of seed microbial communities because seeds are mobile. Microorganisms are recruited from a sequence of species pools: first from the microbiota of the mother tree’s aboveground organs, and later from surrounding microbial sources such as bioaerosols, litter and soil (Barret *et al.*, 2016). The microorganisms that are directly transferred from the vascular system of mother plant to the seedlings through seeds are termed vertically-transmitted (Truyens *et al.*, 2015), while the others are called horizontally-acquired. Vertically-transmitted microorganisms can trigger maternal effects (defined as the causal influence of the maternal genotype or phenotype on the offspring phenotype; Wolf & Wade, 2009) in seed and seedling extended phenotypes (Vivas *et al.*, 2015).

To gain insight into processes of seed microbiome assembly in natural ecosystems, we analyzed the microbial turnover among acorns of sessile oak (*Quercus petraea*) populations growing along elevation gradients. We collected acorns in the canopy and on the ground beneath individual trees, and used Hierarchical Models of Species Communities (HMSC; Ovaskainen *et al.*, 2017) to quantify the ecological processes shaping seed microbial communities at each spatial level (oak population, mother tree and seed) and to generate hypotheses of interactions among microorganisms. We focused on fungal communities as they include pathogens that are detrimental to seed survival (such as *Ciboria batschiana*; Prochazkova *et al.*, 2005). After having described and visualized the fungal communities associated to acorn tissues, we tested the following hypotheses: (**H1**) acorn fungal communities are shaped by environmental filtering and maternal effects, (**H2**) the maternal effects tend to disappear after acorn fall because of horizontal acquisition of fungi from the ground, (**H3**) the response of fungal species to variations in the acorn environment depend on their lifestyle, and (**H4**) biotic interactions among fungal species play a role in the protection of acorns against pathogens.

## Materials and methods

### Sampling design

Acorns were collected on October 21^st^ and 22^nd^ 2015 in four populations of sessile oak located in the Pyrenees Mountains (France). Two populations (Ade and Bager) were at an elevation of ∼400 m a.s.l. and the other two (Gedre-Bas and Gabas) at ∼1200 m a.s.l. The sampling date was chosen as close as possible to the fruiting peak at both elevations (see Caignard *et al.*, 2017 for a complete description of the sampling sites and fruiting phenology). In each site, acorns were collected from three trees randomly selected among the dominant adults. For each tree, we collected four acorns from the canopy using a slingshot and four acorns from the ground beneath the crown (within a distance of 2 m from the trunk). We also collected the biotic microenvironment of canopy acorns, defined as all tree tissues present in a cylinder of 4 cm diameter and 6 cm length around the acorn (including the acorn cupule, the cupules of other acorns, the twig to which the acorn was attached, the leaf petioles and the base of leaves) and the biotic microenvironment of ground acorns, defined as all substrates beneath the acorn within a cylinder of 4 cm diameter and 1 cm depth (including dead oak leaves, dead leaves of other plant species, acorn caps, twigs, pieces of bark, granules of soil, mosses, lichens or herbs). Each sample was collected aseptically, using new plastic gloves and scissors cleaned with 96% ethanol to minimize contamination. Samples were stored in individual plastic vials in a cooler with ice until they could be stored at −80°C.

Ten additional acorns were harvested from five mother trees (two in Gedre-Bas, two in Gabas and one in Ade). For each tree, one acorn was harvested in the canopy and one acorn was harvested on the ground. These acorns were surface-sterilized and dissected to characterize the endophytic fungal communities associated with acorn internal tissues. Surface-sterilization was completed using a three-step process: immersion for 3 min in a 70% ethanol solution, immersion for 2 min in a 3% calcium hypochlorite solution and rinsing with DNAway and sterilized water. After drying the acorns on sterilized filter papers, the fruit walls and the embryos were collected using a sterilized nutcracker and pliers and stored at −80°C.

### DNA extraction, amplification and sequencing of fungal communities

Samples were ground into a homogeneous powder using liquid nitrogen. Mortars and pestles were cleaned using DNAway and autoclaved for 20 min at 121°C between each sample. Approximately 45 mg of powder from each sample was transferred to a microplate under a laminar flow hood. Microplates were stored at −80°C until DNA extraction. Total DNA was extracted using DNeasy Plant Mini Kits (Qiagen, USA) according to the manufacturer’s protocol except that DNA extracts were eluted twice with 50µL of elution buffer (10mM Tris-Cl, 0.5mM EDTA; pH 9.0).

The ITS1 region of the nuclear ribosomal internal transcribed spacer, considered the universal barcode marker for fungi (Schoch *et al.*, 2012), was amplified using the ITS1F (5′-CTTGGTCATTTAGAGGAAGTAA-3′; Gardes & Bruns, 1993) and ITS2 (5′-GCTGCGTTCTTCATCGATGC-3′; White, Bruns, Lee, & Taylor, 1990) primers. The reaction mixture (12.5 µL of final volume) consisted of 1.25 µL of template DNA, 2.5µL of 1µM of each of the forward and reverse primers and 6.25µL of 2X KAPA HiFi HotStart ReadyMix (Kapa Biosystems) containing 0.6 mM of each dNTP, 5 mM MgCl_2_, and 2 units of Kapa Taq DNA polymerase. PCR cycling reactions were conducted on a Veriti 96-well Thermal Cycler (Applied Biosystems) using the following conditions: initial denaturation at 95 °C for 3 min followed by 20 cycles at 98°C for 20 s, 55°C for 45 s, 72°C for 15 s with final extension of 72 °C for 1 min. ITS1 amplification was confirmed by electrophoresis on a 2% agarose gel. Templates that were not successfully amplified using this protocol were amplified again after DNA dilution to 20ng/µL or 10ng/µL. Two marine fungal species (*Yamadazyma barbieri* and *Candida oceani*) were used as positive controls as they were unlikely to be found in our samples. One positive control included DNA of a strain of *C. oceani*, and the other included an equimolar mixture of the DNA of both species. A first negative control was represented by 1 mL of water washes of 4 empty plastic vials opened during the sampling campaign and washed with sterile water. The PCR mix was used as a second negative control.

PCR products were diluted five times in PCR-grade water and used as a DNA template for a second PCR performed using the tailed primers ITS1F_PlaGe (5’-CTTTCCCTACACGACGCTCTTCCGATCTCTTGGTCATTTAGAGGAAGTAA-3’) and ITS2_PlaGe (5’-GGAGTTCAGACGTGTGCTCTTCCGATCTGCTGCGTTCTTCATCGATGC-3’) designed by the Get-PlaGe sequencing facility (Toulouse, France). The second PCR was performed twice (once in 12.5 µL and once in 25µL of final volume) using the same reaction mixture as the first PCR. The PCR conditions were as follows: initial denaturation at 95 °C for 3 min followed by 10 cycles at 98°C for 20 s, 60°C for 15 s, 72°C for 15 s and a final extension step of 72°C for 1 min. PCR products were purified (CleanPCR, MokaScience), multiplex identifiers and sequencing adapters were added, and library sequencing on an Illumina MiSeq platform (v3 chemistry, 2×250 bp) and sequence demultiplexing (with exact index search) were performed at the Get-PlaGe sequencing facility (Toulouse, France).

### Bioinformatic analyses

Paired-end sequences were joined using PEAR v0.9.10 (Zhang *et al.*, 2014). Only pairs with a minimum overlap of 50 bp and without any uncalled bases were kept. Assembled sequences were filtered using DADA2 v1.4.0 (Callahan *et al.*, 2016). Only sequences with less than one expected error and longer than 100 bp were retained in the dataset. Amplicon sequence variants (ASVs) were inferred using DADA2 and chimeric sequences were removed using the consensus method of the *removeBimeras* function. Taxonomic assignments were performed using the RDP classifier (Wang *et al.*, 2007) implemented in DADA2 and trained with the UNITE database 7.2 (UNITE Community 2017), with an 80% confidence threshold. The ASV table was then imported in R using the phyloseq package v1.26.0 (McMurdie & Holmes, 2013) and filtered. Only ASVs assigned to a fungal phylum were kept. Positive and negative controls were used to remove contaminants (Galan *et al.*, 2016). The cross-contamination threshold (T_CC_) was defined as the maximal number of sequences of each ASV found in negative or positive control samples. The false-assignment threshold (T_FA_) was defined as the highest sequence count of a positive control strain in a non-control sample, divided by the total number of sequences of the strain in the whole run and multiplied by the total number of sequences of each ASV. ASVs were removed from all samples where they harbored fewer sequences than either threshold (T_FA_ or T_CC_).

### SNP genotyping and maternity exclusion analyses

Acorns collected on the ground were genotyped to confirm that they belonged to the mother tree above them. Genotyping was performed using 39 polymorphic single nucleotide polymorphism (SNP) markers (Gerzabek *et al.*, 2017). DNA was diluted to a final concentration of 15 to 20 ng/µL and sequenced using the iPLEX Gold Genotyping kit (Agena, San Diego, CA, USA) at the Genome Transcriptome Facility of Bordeaux (PGTB, Bordeaux, France) according to the manufacturer’s instructions (see Chancerel *et al.*, 2013 for more details). Two samples of aboveground tissues of each tree were genotyped and compared to estimate the typing error rate of false calls during genotyping. Loci with poor performance during the clustering procedure (call rates <60%) were excluded, resulting in a final set of 28 loci. Acorn genotypes were compared to the genotype of their putative mother tree. Considering the low error rate of these SNPs (Gerzabek *et al.*, 2017), we took a deliberately conservative approach and assumed that if tree and acorn shared no alleles for at least one locus, the mother-offspring relationship was not confirmed and the acorn sample was removed from the dataset.

### Confocal microscopy

Sixteen additional acorns were collected in autumn 2017 from the ground of the oak forest of Bellebat (44°43’36.4”N 0°13’22.5”W, Southwest of France) to visualize fungal colonization outside and inside acorns. Acorns were cut in half with secators and fixed overnight at 4°C in a paraformaldehyde solution (4% w/v in PBS, pH 7.2). Samples were then rinsed three times with PBS, immersed in 15 mL PBS containing 50 μg ml^-1^ of wheat germ agglutinin (WGA)-AlexaFluor488 conjugate (Life Technologies, USA), incubated 2 hours at 37°C, and rinsed again three times with PBS. The samples were observed under a confocal microscope (Olympus Fluoview FV1000 with multiline laser FV5-LAMAR-2 and HeNe(G) laser FV10-LAHEG230-2). Pictures were taken with X, Y, Z coordinates at 405, 488, 594 nm and with 10X, 20X, 40X or 60X objectives. Images were merged (RGB) using Image J software (Schneider *et al.* 2012). Pictures were created using Z Project Stacks (Campisano *et al.* 2014), then cropped, and the light/contrast balance was improved (Glassner *et al.* 2015). Images presented in this publication correspond to the average colonization level observed.

### Statistical analyses

#### Comparison of acorn fungal communities among mother trees and environments

To test hypotheses H1 and H2, we analyzed the effects of mother tree, local environment (oak population), and microenvironment (acorn position in the canopy *versus* on the ground) on acorn fungal community richness and composition. Fungal richness was defined as the total number of ASVs per acorn sample and was modelled using generalized linear mixed models (GLMMs) with a negative binomial distribution and a log-link function. The first model had oak population, acorn position and their interaction as fixed effects and the mother tree as a random effect. We then tested the effect of the mother tree on fungal richness for acorns in the canopy and acorns on the ground separately. The two models had the mother tree as fixed effect and the population as random effect. The natural logarithm of the total number of sequences per sample (sequencing depth) was introduced as an offset in all models.

We then analyzed the effects of the three same factors on fungal community composition by using permutational multivariate analyses of variance (PERMANOVAs) with 9999 permutations. Compositional dissimilarities among acorn samples were estimated using quantitative and binary versions of the Jaccard index (Jaccard, 1901) and visualized with principal coordinate analyses (PCoA). We first tested the effects of oak population, acorn position and their interaction on compositional dissimilarities among samples, by constraining or not constraining the permutations by elevation. We then assessed the effect of mother tree on community composition for acorns in the canopy and on the ground separately. The two models had the mother tree as a fixed effect and permutations were constrained by population. The natural logarithm of the total number of sequences per sample (sequencing depth) was introduced as the first effect in all models.

In addition, we investigated whether changes in fungal community composition after acorn fall were due to either the substitution of canopy-associated fungal species by ground-associated fungal species or gain of ground-associated fungal species without loss of canopy-associated fungal species, by partitioning Jaccard binary dissimilarities among acorns of the same mother tree using the betapart package v1.5.1 (Baselga *et al.*, 2018). The proportion of fungal species of acorns on the ground also found in acorns in the canopy was calculated for each mother tree.

#### Quantification of maternal effects, environmental filtering and biotic interactions

Hierarchical Models of Species Communities (HMSC; Ovaskainen *et al.*, 2017) were then used to quantify processes of community assembly. The models assumed that variation in fungal community composition among acorns (i.e. the **Y** matrix in the HMSC framework) was accounted for by four ecological predictors (*Mother mycobiota, Microenv mycobiota, Acorn position* and *Site elevation*) introduced as fixed effects in the **X** matrix. The **Y** matrix represented ASV sequence counts in all acorn samples, out of which we included only the ASVs that were present in five or more acorns. *Mother mycobiota* and *Microenv mycobiota* represented fungal communities in the canopy of mothers trees and in the microenvironment of acorns, respectively. They were ASV-specific predictors and thus the **X** matrix was different for every ASV of the **Y** matrix. *Mother mycobiota* was calculated for all acorn samples as the average relative abundance of the focal ASV in the twigs and leaves of mother trees, and was included at the tree level to model vertical transmission of fungi from the mother tree to its acorns. *Microenv mycobiota* was calculated as the residuals of the regression of the relative abundance of the focal ASV in the microenvironment of each acorn over its relative abundance in the canopy of the mother tree, and was included at the sample level to model horizontal acquisition of fungi from the materials surrounding each acorn. *Site elevation* and *Acorn position* represented filtering of fungal communities by climate and microclimate, respectively. *Site elevation* was included at the site level to model selection exerted by site-level abiotic factors, such as average air temperature, on acorn fungal communities. *Acorn position* (canopy vs. ground) was included at the sample level to model selection exerted by microclimate, such as higher humidity on the ground, on acorn fungal communities. *Mother mycobiota* and *Microenv mycobiota* were included in interaction with *Acorn position* to test the hypothesis that their contribution to fungal communities differ between acorns in the canopy and acorns on the ground (H2). We also introduced the log-transformed sequencing depth of each sample (*Sequencing depth*) as a fixed effect in the **X** matrix, to take into account methodological biases influencing ASV sequence counts. Random effects at each hierarchical level (oak population, mother tree and acorn sample) were also introduced to model variations in ASV sequence counts that can neither be attributed to the four ecological predictors nor sequencing depth. In addition, we tested the hypothesis that fungal lifestyle modulates fungal ASV responses to environmental variations (H3), by including the trophic mode (saprotroph, plant pathogen or other) and the degree of specialization toward acorns of each ASV in the **T** matrix. The putative trophic mode of each ASV was determined using the FUNGuild database (Nguyen *et al.*, 2016). Their degree of specialization toward acorns was defined as the log-ratio of ASV relative abundance in acorns *versus* other sample types (i.e. branches, leaves, litter and upper soil), calculated using DESeq2 (Love *et al.*, 2014). To account for the zero-inflated nature of the data, we applied a hurdle modelling approach. We first fitted a probit model on ASV presence-absence data and then fitted a linear model on sequence count data conditional on presence, in which counts were log-transformed and scaled to zero mean and unit variance for each ASV and absences masked as non-available data. We fitted both models with default prior distributions (Ovaskainen *et al.*, 2017). For each of the four MCMC chains, we sampled the posterior for 1,500,000 iterations, out of which we excluded the first 500,000 as burn-in and thinned the remaining iterations by 1000, thus producing a total of 4,000 posterior samples. We examined MCMC convergence through the distributions of potential scale reduction factors (PSRF) of the model parameters. To examine model fit, we applied a two-fold cross validation across the samples and evaluated predictive performance by AUC for the presence-absence model and *R*^2^ for the linear model. Finally, residual correlations among fungal ASV sequence counts at the acorn level were interpreted as hypothetical biotic interactions among fungal strains (see Ovaskainen *et al.*, 2017). We examined associations among ASVs assigned at the species level to test hypothesis H4.

## Results

All acorn tissues, including the fruit wall, seed coat, and embryo were colonized by fungi, with a dense colonization under the endocarp (Fig. 1A-E). Ascomycota represented 91.1% and 89.4% of the sequences of canopy and ground acorns, respectively. *Dothideomycetes, Leotiomycetes* and *Sordariomycetes* were the three main classes of ascomycetes present (Fig. S1). Among the ten most abundant species associated to whole acorns, five are known as plant pathogens (*Gnomoniopsis paraclavulata, Taphrina carpini, Epicoccum nigrum, Mycosphaerella tassiana* and *Polyscytalum algarvense*) and two are known as antagonists of other microorganisms (*Cladosporium delicatulum, Cylindrium elongatum*) (Table 1; Tables S2 and S3 for all ASVs and subset of ASVs used in HMSC models, respectively). The ubiquitous fungi *Curvibasidium cygneicolum* and *Epicoccum nigrum* were dominant in the internal tissues of acorns, including the embryo (Table S1).

**Table 1.** Most abundant fungal species associated with seeds of sessile oak and their microenvironment. Seeds were either collected in the tree canopy or on the ground. Materials from the seed microenvironment (twigs and leaves, or litter and upper soil) were also collected. The fungal community of all 4 sample types was analyzed using a metabarcoding approach. Only Amplicon sequence variants (ASV) assigned to Ascomycota (A) or Basidiomycota (B) with the UNITE database were kept. Average relative abundances of all ASVs were computed for each sample type, after merging ASVs assigned to the same fungal species. Only ASVs identified at the species level are shown in the table.

| Fungal species | Average relative abundance per sample type (%) |  |  |  |  | Lifestyle | Reference(s) |
| --- | --- | --- | --- | --- | --- | --- | --- |
|  | All seeds | Seeds in the canopy | Canopy samples (twigs, leaves) | Seeds on the ground | Ground samples (litter, upper soil) |  |  |
| *Gnomoniopsis paraclavulata* (A) | 13.4 | 5.3 | 1.5 | 21.9 | 0 | Pathogen isolated in leaves, buds, cupules and shoots of *Castanea sativa*. Commonly isolated from overwintered leaves of *Quercus* sp. | Tosi *et al.* (2014) ; Sogonov *et al.* (2008) |
| *Stromatoseptoria castaneicola* (A) | 2.7 | 5.2 | 0.9 | 0 | 0.2 | Causes leaf spots on *Castanea sativa*. | Quaedvlieg *et al.* (2013) |
| *Taphrina carpini* (A) | 2.4 | 3.4 | 0.7 | 1.4 | 0.3 | Common pathogen encountered on *Fagaceae* leaves. | Bacigálová (1991); Inácio *et al.* (2004); Cordier *et al.* (2012) |
| *Epicoccum nigrum* (A) | 2.3 | 0.4 | 0.1 | 4.2 | 0.2 | Ubiquitous fungus found in soil, leaves and seeds described as primary saprotroph and plant pathogen. | Ahumada-Rudolph *et al.* (2014) |
| *Mycosphaerella tassiana* (A) | 1.7 | 1.5 | 0.1 | 1.9 | 0.3 | Common pathogen found in the phyllosphere including that of oak. | Schubert *et al.* (2007); Jakuschkin *et al.* (2016) |
| *Curvibasidium cygneicollum* (B) | 1.5 | 1.2 | 0.2 | 1.9 | 0.1 | Endophyte of fruits, leaves, trunk and soil behaving as a phytopathogen or a mycoparasite. The species is insensitive to the mycocins produce by *Filobasidium* and *Cystofilobasidium*. | Sampaio *et al.* (2004); Mašínová *et al.* (2017) ; Sampaio *et al.* (2004) |
| *Cylindrium elongatum* (A) | 1.5 | 1.4 | 0.1 | 1.5 | 4.3 | Bacterial and fungal antagonist found on oak leaves. | Reyes-Estebanez (2011) ; Duarte *et al.* (2015) |
| *Polyscytalum algarvense* (A) | 1.5 | 1.9 | 0.1 | 1 | 5.3 | Necrotroph fungi found on *Eucalyptus* leaves. | Cheewangkoon *et al.* (2009); Crous (2018) |
| *Fusarium pseudensiforme* (A) | 1.3 | 1.4 | 0 | 1.2 | 0 | Found on bark of trees. | Nalim *et al.* (2011) |
| *Cladosporium delicatulum* (A) | 1.3 | 1.3 | 0.2 | 1.3 | 0.3 | Found in cereal seeds, mycoparasite of *Taphrina* spp and *Magnaporthe oryzae* | Amanelah Baharvandi & Zafari (2015) ; Chaibub *et al.* (2016) |

**Figure 1.**
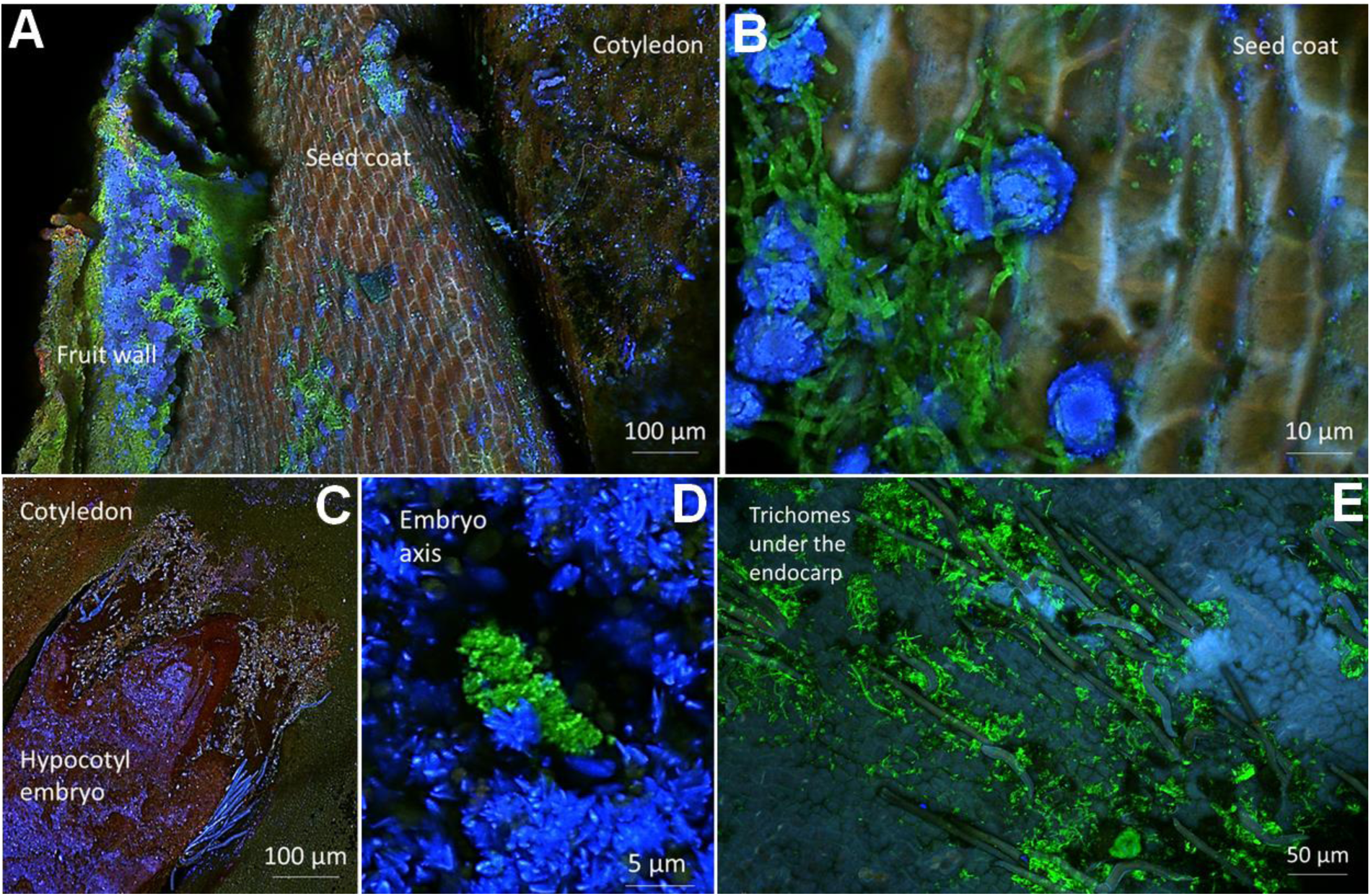
Fungi in cross sections of seeds of sessile oak collected on the ground. Fungi (green fluorescent) were revealed by confocal microscopy and WGA-ALEXA fluor488 staining. (A) Fruit wall and seed coat. (B) Zoom-in view of panel A. (C) Embryo and cotyledon. (D) Zoom-in view of panel C. (E) Internal surface of the endocarp.

### Acorn fungal communities are shaped by environmental filtering and maternal effects

Acorn fungal community richness (Table 2) and composition (Tables 3 and S2) differed significantly among oak populations along elevation gradients, and among mother trees within the same population, confirming the hypothesis that both environmental filtering and maternal effects shape acorn fungal communities (H1). PCoAs suggested that fungal communities were more similar between populations at the same elevation (Fig. 2). However, the population effect remained significant when PERMANOVAs were constrained by elevation (Pseudo-*F* = 1.24, *P* < 0.01), implying that selection exerted by abiotic conditions along the gradients was not the only process triggering variation in fungal community composition among oak populations.

**Table 2.** Generalized linear mixed-effects models (GLMM) of fungal community richness of seeds of sessile oak. Richness was defined as the number of amplicon sequence variants (ASV) per sample. The total number of sequences per sample (sequencing depth, SD) was introduced as an offset in all models. The effects of tree population (T), seed position (canopy *versus* ground, P) and their interaction were tested on the whole seed dataset while the effect of mother tree was tested separately on canopy seeds and ground seeds.

|  | d.f. | chi-square | P-value |
| --- | --- | --- | --- |
| **All seeds** |  |  |  |
| Tree population (T) | 3 | 12.5 | **<.01** |
| Seed position (P) | 1 | 12.6 | **<.001** |
| T × P | 3 | 5.74 | 0.12 |
| **Seeds in the canopy** |  |  |  |
| Mother Tree | 10 | 30.6 | **<.01** |
| **Seeds on the ground** |  |  |  |
| Mother Tree | 11 | 28.3 | **<.01** |

**Table 3.** Permutational multivariate analyses of variance (PERMANOVA) of compositional dissimilarities among fungal communities of seeds of sessile oak. Dissimilarities among seeds were estimated using the binary Jaccard distance. The total number of sequences per sample (sequencing depth, SD) was log-transformed and introduced as the first explanatory variable in all models. The effects of tree population (T), seed position (canopy *versus* ground, P) and their interaction were tested on the whole seed dataset while the effect of mother tree was tested separately on canopy seeds and ground seeds.

|  | d.f. | F-value | P-value | R<sup>2</sup> |
| --- | --- | --- | --- | --- |
| **All seeds** |  |  |  |  |
| Log(SD) | 1 | 1.3 | <.01 | 0.01585 |
| Tree population (T) | 3 | 2.4 | <.001 | 0.08711 |
| Seed position (P) | 1 | 1.8 | <.001 | 0.02226 |
| S x P | 3 | 1.2 | <.001 | 0.04494 |
| **Seeds in the canopy** |  |  |  |  |
| Log(SD) | 1 | 1.3 | 0.191 | 0.03052 |
| Mother tree | 10 | 1.4 | **0.010** | 0.32742 |
| **Seeds on the ground** |  |  |  |  |
| Log(SD) | 1 | 1.1 | 0.319 | 0.02834 |
| Mother tree | 11 | 1.2 | **0.025** | 0.34149 |

**Figure 2.**
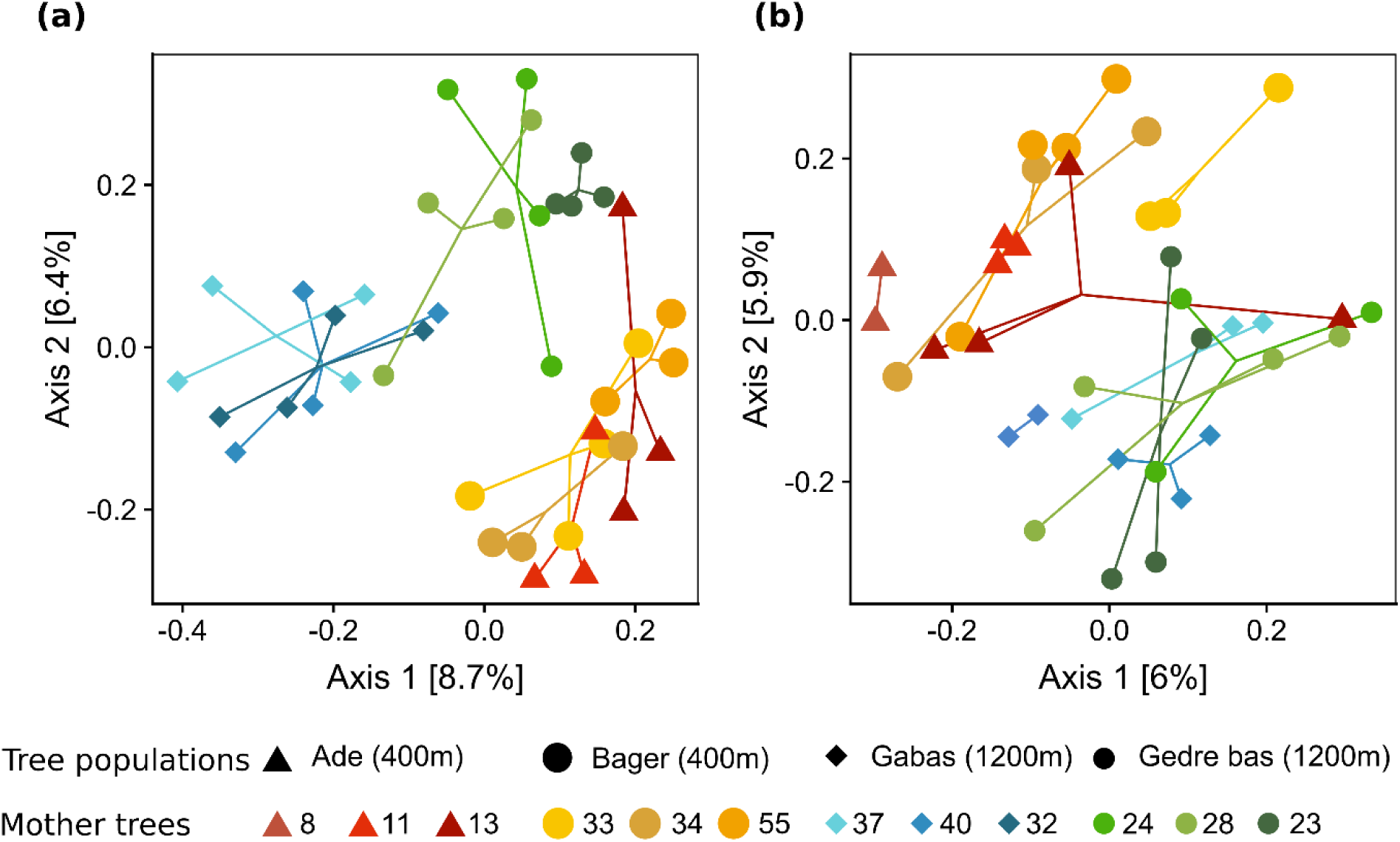
Compositional dissimilarities among fungal communities of seeds collected (A) in the canopy and (B) on the ground in four populations of sessile oak. Dissimilarities among seeds were estimated using binary Jaccard distance and represented with a PCoA plot. Fungal community composition differed significantly among tree populations and among mother trees (Table 3).

PCoAs and HMSC models gave different estimates of the relative contribution of environmental filtering and maternals effects. PCoAs suggested that mother tree identity had a lower influence on fungal community composition than elevation (Fig. 2), whereas HMSC models indicated the opposite trend (Table 4). The model of ASV presence-absence explained (in units of AUC, averaged over the ASVs) 80% of the variation for the model fitted to all data and 69% of the variation based on the two-fold cross-validation approach. According to this model, elevation was a minor direct driver of fungal community composition (only 2% of the explained variance). In contrast, the average relative abundance of a fungal ASV in the tissues of a mother tree (*Mother mycobiota*) was the second most important predictor of the occurrence of this ASV in an acorn from this tree, in interaction with the acorn position (39% of the explained variance). *Mother mycobiota* was the unique predictor of occurrence for several ASVs belonging to the orders *Helotiales, Venturiales* and *Xylariales* (Fig. S2). A similar ranking of the predictors was obtained for the model of ASV sequence counts (Table 4), except that sequencing depth and random effects had a much larger influence on ASV sequence counts than on ASV presence-absence (17% *versus* 1% and 21% *versus* 6%, respectively). This model only explained 34% of the variation for the model fitted to all data and 18% of the variation based on the two-fold cross-validation approach.

**Table 4.**
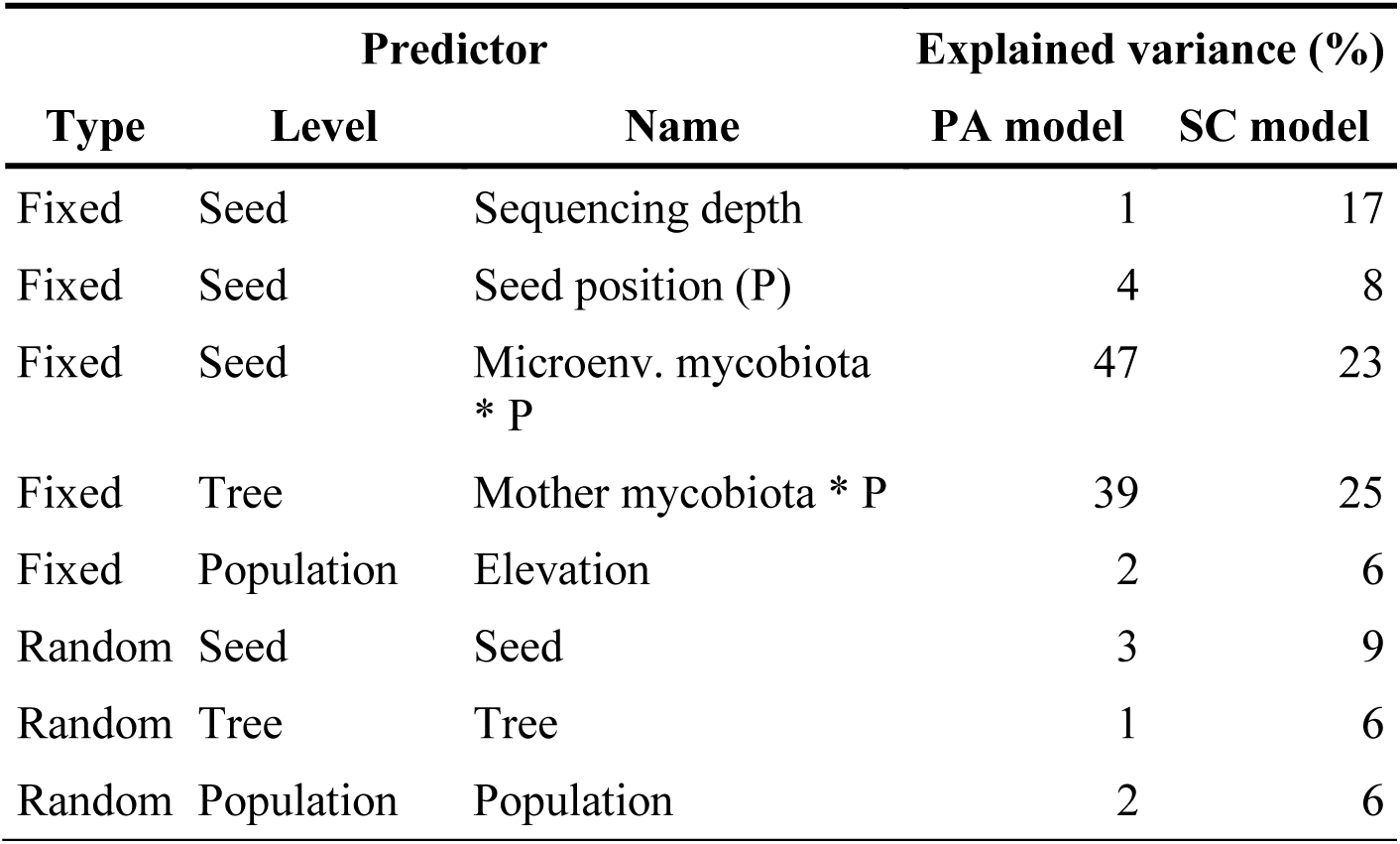
Partitioning of the variance in fungal community composition of seeds of sessile oak. Four fixed effects were included in the HMSC models to explain variations in the presence-absence (PA) or the sequence count (SC) of a focal fungal ASV among seeds: *Sequencing depth* (total number of sequences per sample), *Seed position* (canopy or ground), *Microenv. mycobiota* (relative abundance of the focal ASV in the seed biotic microenvironment), *Mother mycobiota* (average relative abundance of the focal ASV in the mother tree aboveground tissues), and *Elevation*. Random effects were included at each spatial scale (seed, tree and population). Results of variance partitioning are given as percentages (%) of total explained variance.

### Maternal effects persist after acorn fall despite horizontal acquisition of fungi from the ground

Fungal community richness (Table 2) and composition (Table 3 and S2) differed significantly between acorns in the canopy and acorns on the ground. For instance, *Gnomoniopsis paraclavulata* was four times more abundant in acorns on the ground than in acorns in the canopy, while *Stromatospheria castaneicola* was only present in canopy acorns (Table 1). Fungal richness increased and composition shifted toward that of ground materials after acorn fall (Fig. 3), confirming the horizontal acquisition of fungi from the ground. Partitioning of Jaccard beta-diversity indicated that these temporal changes in community composition were mainly driven by turnover (replacement of fungal species rather than net gains or losses in species number; Table S4). HMSC models confirmed the large influence of horizontal transmission on acorn fungal communities. The relative abundance of a fungal ASV in the microenvironment of an acorn (*Microenv mycobiota*), in interaction with the acorn position, was generally the best predictor of ASV occurrence (47% of the explained variance), especially for the *Capnodiales, Dothideales* and *Taphrinales* orders (Fig. S2).

**Figure 3.**
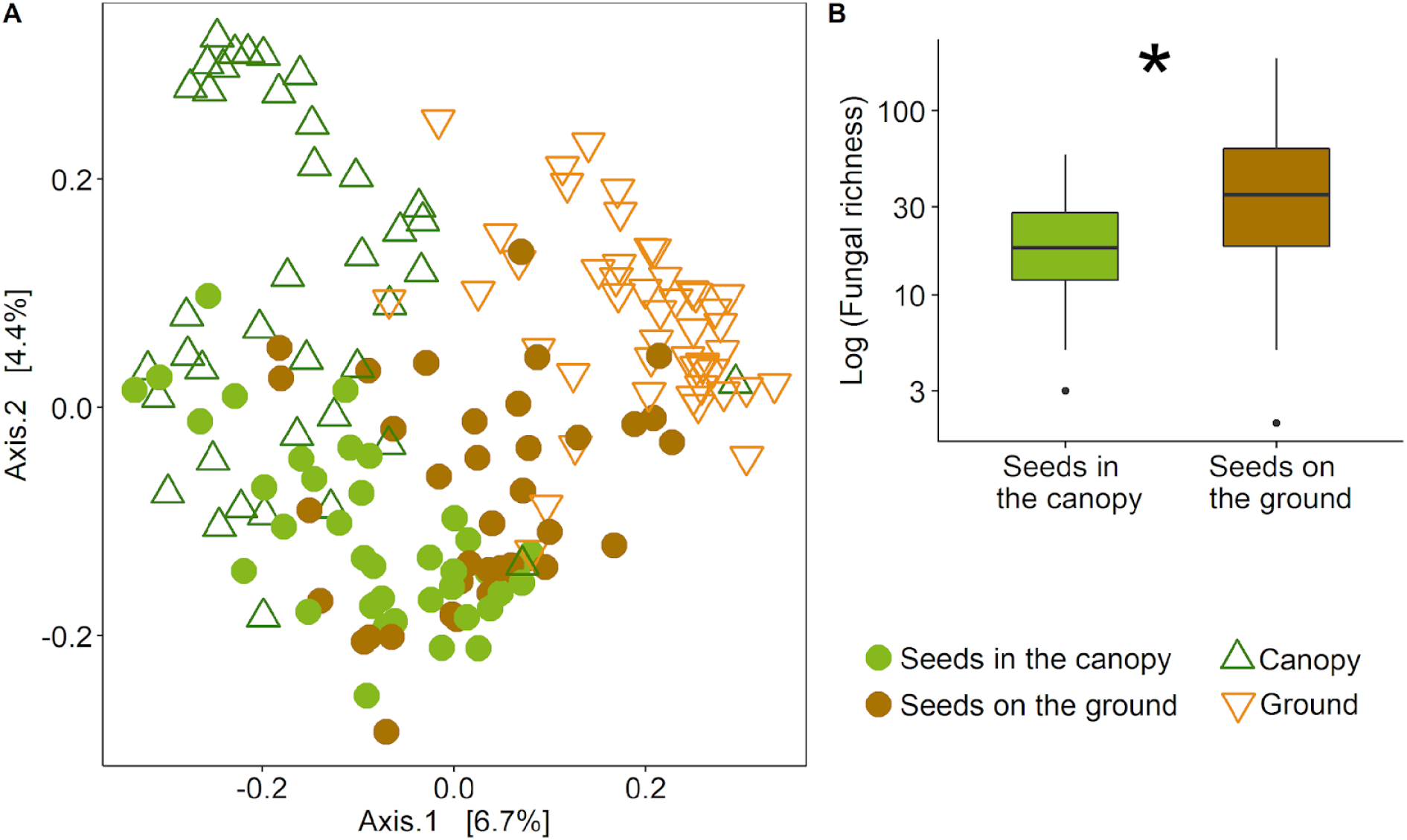
Fungal community composition and richness of seeds of sessile oak collected in the canopy and on the ground. (A) PCoA plot of compositional dissimilarities among fungal communities associated to seeds, canopy (leaves and twigs) and ground materials (litter and upper soil). Dissimilarities among samples were estimated using binary Jaccard distance. Fungal community composition differed significantly among the four sample types (PERMANOVA; F=4.3, *P*<0.001). (B) Richness (log-transformed) of seed fungal communities, defined as the number of ASVs per sample. Richness was significantly higher in seeds on the ground (Table 2).

**Figure 4.**
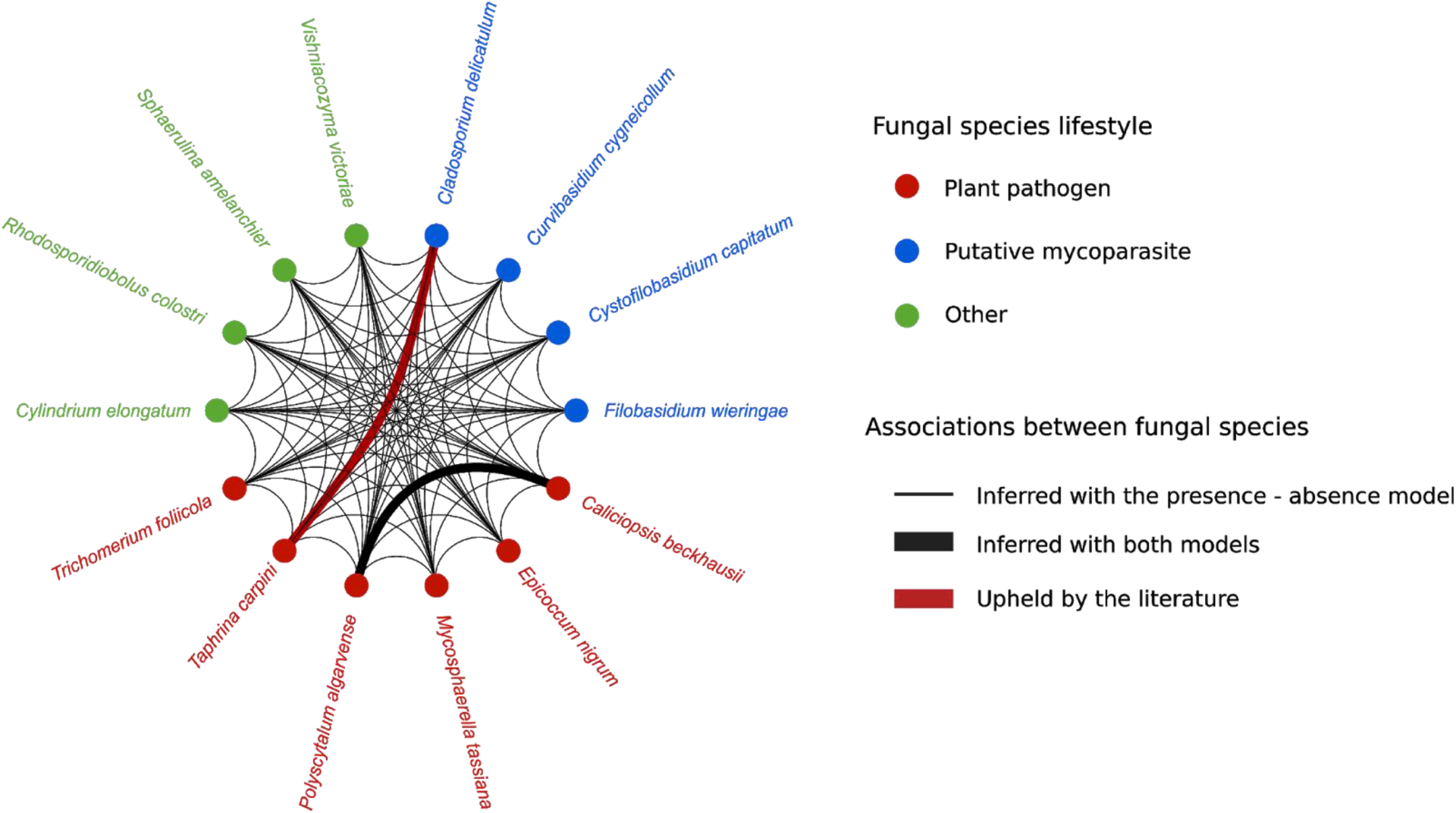
Network of fungal associations estimated by HMSC models. Associations were all positive. Network nodes correspond to fungal ASVs assigned to the species level and their color correspond to putative functions according to FUNGuild and the literature (Table S6). Network links indicate associations with at least 95% posterior probability estimated by the presence-absence model (thin black plain line) or by both the presence-absence model and the sequence count model (thick black plain line). The positive association between *Taphrina carpini* and *Cladosporium delicatulum* is indicated in red because it is described in the literature as a host-parasite interaction (Table S6).

Despite the turnover in fungal community composition after acorn fall and in contrast with our hypothesis (H2), the maternal effects were significant for acorns on the ground (Table 3), indicating that some maternal species were retained after acorn fall. Overall, acorns on the ground shared 10 to 40% of their fungal community with acorns in the canopy of the same mother tree, and 21 to 50% with mother tree tissues (Table S5). On average, 38% of fungal ASVs of acorns on the ground were present in both acorns in the canopy and mother tree tissues. Fungal species most often retained after acorn fall were *Taphrina* sp., *Cladosporium delicatulum*, a mycoparasite of *Taphrina* sp. (Baharvandi & Zafari, 2015), and the ubiquitous *Epicoccum nigrum*.

### The response of fungal species to variations in the acorn environment depend on their lifestyle

HMSC models showed that fungal lifestyle influenced the response of acorn fungal communities to environmental variations, in accordance with our hypothesis (H3). High elevation selected for saprotroph species and seed specialists, increasing their proportion in acorn fungal communities (Table S3). Acorn fall favored pathogen species and saprotrophs. Their proportion increased after acorn fall while their abundance (conditional on presence) was not altered (Table S3). These findings suggest that vertically-transmitted pathogens did not increase in abundance after acorn fall, and that acorns were colonized by pathogen and saprotroph species of the ground.

### Biotic interactions among fungal species might play a role in the protection of acorns against pathogens

Fungal colonization was very dense on acorn external surfaces (Fig. 1A) but also on internal surface of the fruit wall (Fig. 1E), indicating that fungal colonizers might enter into contact and compete for space and eventually resources. In contrast to this expectation, residual co-occurrence patterns of HMSC models at the acorn level revealed only positive associations between fungal ASVs. Among fungal ASVs associated with each other, 14 could be assigned at the species level (Fig. 5). Six of them are described as plant pathogens and two of them, *Mycosphaerella tassiana* and *Taphrina carpini*, have already been found in association with oak (Table S6). Four of the species are described as mycoparasites, including *Cladosporium delicatulum*. The positive association between *Taphrina carpini* and *Cladosporium delicatulum* (Fig. 5), which are both found in the embryo and the fruit wall (Table S3), might therefore represent a vertically-transmitted host-parasite interaction. This interaction might play a role in oak disease regulation, in accordance with our hypothesis (H4).

## Discussion

Seed microbial communities of trees have received little attention so far despite their potential influence on forest dynamics and evolution. In this study, we compared fungal communities of seeds among four populations of sessile oak (*Quercus petraea*), a dominant tree species of deciduous temperate European forests (McShea *et al.*, 2006). The populations were located along elevation gradients in the Pyrenees Mountains (France). They differed by up to 800 m in elevation. Our analyses revealed that the richness and composition of seed fungal communities differed significantly among oak populations along gradients, suggesting that abiotic filtering was a major driver of community assembly. However, Hierarchical Models of Species Communities (HMSC; Ovaskainen et al. 2017) revealed that the direct influence of elevation on seed fungal communities was small. Elevation accounted for only 2% of the explained variance in fungal species presence-absence, and 6% of the variance in species abundance. The apparently large effects of elevation suggested by multivariate analyses might therefore be indirect effects. HMSC models estimated that fungal community composition in the microenvironment of each seed (i.e. twigs and leaves surrounding seeds in the canopy, or ground materials surrounding dropped seeds), explained 47% of the variance in species presence-absence, and was by far the strongest driver of seed fungal communities. These findings suggest that environmental filtering act directly on the fungal communities of leaves, litter and soil (as shown by Cordier et al., 2012; Coince et al., 2014; Vacher et al., 2016), to then indirectly shape seed communities.

Seed fungal communities also differed among mother trees within the same population and these differences remained significant after seed fall. Such maternal effects have already been found for the abundance of some fungal genera in seeds (Johnston-Monje & Raizada, 2011; Pinciroli *et al.*, 2013). Here we showed that maternal effects extend to the whole fungal community of seeds through to seed fall. HMSC indicated that the fungal community composition of mother tree tissues (twigs and leaves) had a major, direct influence on that of seeds. It accounted for 39% of the explained variance in fungal species presence-absence, and 25% of the variance in species abundance. Recent findings by Vivas *et al.*, (2017) on *Eucalyptus* trees indicate that these maternal effects can persist in seedlings and influence their growth and resistance to pathogens. Together, these results suggest that maternal effects in seed and seedling extended phenotypes could be a major driver of forest regeneration success.

In addition, our confocal microscopy analyses revealed, for the first time, the presence of fungal aggregates within embryos of acorns of sessile oak, as well as a dense fungal colonization on internal surfaces of fruit walls. These endophytic fungal populations contained foliar pathogens of *Fagaceae* tree species, such as *Mycosphaerella tassiana* and *Taphrina carpini* (Schubert *et al.*, 2007; Bacigálová, 1991), ubiquitous fungal species, such as *Epicoccum nigrum* (Andrews & Harris, 2000), and endophytic yeasts previously described in other fruits, such as *Curvibasidium cygneicollum* (Sampaio *et al.*, 2004; Mašínová *et al.*, 2017). Network inference analyses revealed a positive association between the foliar pathogen *Taphrina carpini* and the mycoparasite *Cladosporium delicatulum*, suggesting that mother trees do not only transmit pathogens but also pathogen antagonists. Our results hence confirm that fungal pathogens use seeds for their own dispersion, and that the fungal pathogen’s parasites can follow them using the same dispersion mode (Ewald, 1989; Feldman *et al.*, 2008). Unraveling the genetic architecture of these tripartite interactions, involving tree seedlings, seed-borne pathogens and their hyperparasites, could improve our understanding of forest ecosystem dynamics and evolution.

Finally, our analyses confirmed that seed fall corresponds to a major transition in seed fungal communities. Our results showed that fungal community richness significantly increased and that composition shifted toward that of ground materials after seed fall, confirming that seeds on the ground are rapidly colonized by the species present in the surrounding microenvironment (Crist, 2009, Qin *et al.*, 2016; Truyens *et al.*, 2015; Klaedtke *et al.*, 2016). For instance, the species *Gnomoniopsis paraclavulata*, that was previously found in association with oak litter (U’Ren *et al.*, 2016), drastically increased in abundance after acorn fall. Our analyses also suggested that seed fall triggers a replacement of canopy-inherited species by ground-derived species, rather than an addition of species associated to ground materials. This replacement was however only partial. Seeds on the ground shared up to 50% of their fungal colonizers with the twigs and leaves of their mother tree. The mechanisms of community filtration during vertical transmission (Vannier *et al.*, 2018) will have to be investigated in future studies.

## Conclusions

Our study revealed that acorns of sessile oak harbor diversified fungal communities in their internal tissues, including the embryo, and on their surfaces. These communities were shaped by maternal effects, environmental filtering and biotic interactions. Maternal effects persisted after seed fall, despite seed colonization by soil and litter fungi. Environmental filtering did not shape directly seed fungal communities, but rather influenced communities in the microenvironments surrounding seeds. Biotic interactions included several host-parasite interactions between tree pathogens and their antagonists, one of which was likely to be vertically-transmitted. Future research will have to investigate the maternal and environmental drivers of the rate of vertical transmission of microorganisms (e.g. Cavazos *et al.*, 2018; Sneck *et al.*, 2017; Leff *et al.*, 2017), and assess the role of these microorganisms on seed survival and germination, seedling growth and health (e.g. Vivas *et al.*, 2017; Leroy *et al.*, 2019), and ultimately tree fitness. The influence of vertically-transmitted microorganisms on seed and seedling secondary metabolites (Chen *et al.*, 2018; Shazad *et al.*, 2018), and their cascading effects on tree biotic interactions (e.g. Peris *et al.*, 2018) will also have to be investigated. Previous research on oak trees showed for instance a significant relationship between a fungus-like pathogen associated with acorns and the abundance of several oak-dependant bird species, including seed dispersers (Monahan & Koenig, 2006). A combination of germination experiments in controlled conditions (e.g. Leroy *et al.*, 2019), seed microbiome analyses in common gardens (Vivas *et al.*, 2015), and seed microorganisms manipulation across environments (Gundel *et al.*, 2017) will be required to integrate seed microbial ecology into predictive models of forest dynamics and evolution.

## Acknowledgements

We thank Albin Lobo (University of Copenhagen, Denmark) and Jean-Marc Louvet (Biogeco, INRA-University of Bordeaux, France) for their assistance during sampling campaigns. We are also very grateful to Martine-Martin Clotté, Céline Lalanne, Patrick Leger, Adline Delcamp (Biogeco, INRA-University of Bordeaux, France) and Jessica Vallance (SAVE, INRA-Bordeaux Sciences Agro, France) for providing materials and reagents for processing the samples and advice on molecular biology protocols. We thank Samuel Pennarun and all members of the Genotoul sequencing facility (Get-PlaGe, Toulouse, France) for performing MiSeq sequencing, and the Genotoul bioinformatics facility (Bioinfo Genotoul, Toulouse, France) for providing computing and storage resources. We also thank Erwan Guichoux and all members of the Genome Transcriptome facility of Bordeaux (PGTB, Cestas, France) for performing the SNP genotyping. Thanks to Franck Deniel and Gaëtan Burgaud (ESIAB, Université de Bretagne Occidentale, France) for giving us marine fungal strains used as positive controls. We would like to acknowledge George Washington University for its contribution to the funding of molecular biology analyses. We also thank the INRA Meta-Omics of Microbial Ecosystems (MEM) metaprogramme (Learn-biocontrol project), the INRA Ecosystem Services (EcoServ) metaprogramme (IBISC project), the LABEX COTE (MICROMIC project; ANR-10-LABX-45), the LABEX CEBA (ANR-10-LABX-25-01) and the Aquitaine Region (Athene project, n°2016-1R20301-00007218) for additional financial support throughout the project. TF’s PhD grant was funded by the University of Bordeaux. OO was supported by the Academy of Finland (grant 309581), the Erkko foundation, and the Research Council of Norway (SFF-III grant 223257).

## Author contributions

TF participated in the sampling campaigns, processed all samples, extracted and amplified DNA, performed the statistical analyses, interpreted the results and wrote the first draft of the manuscript. CP helped process the samples, performed the bioinformatic analyses and provided statistical analysis scripts. AEZ assisted with supervision and writing. OO conducted the HMSC analysis and helped interpreting the results. TC organized the sampling campaigns and provided environmental data. MB provided advice on molecular biology protocols, bioinformatics and statistical analyses. SC took the microscopy pictures and interpreted the results. AH provided advice and analysis tools for SNP genotyping. SD, AEZ and CV had the original idea for the study. CV participated in the sampling campaigns, coordinated and supervised all stages of the work, and made a major contribution to the writing of the manuscript. All authors revised the manuscript and approved the final version.

## Data availability

All raw sequences obtained from the sequencing of acorns and their biotic microenvironment are available from the National Center for Biotechnology Information Sequence Read Archive (http://www.ncbi.nlm.nih.gov/sra) under the accession number <u>PRJNA551388</u>. The code and ASV tables are available as an archive at https://doi.org/10.15454/SM6OCR.

## Supplementary Materials

**Table S1.**
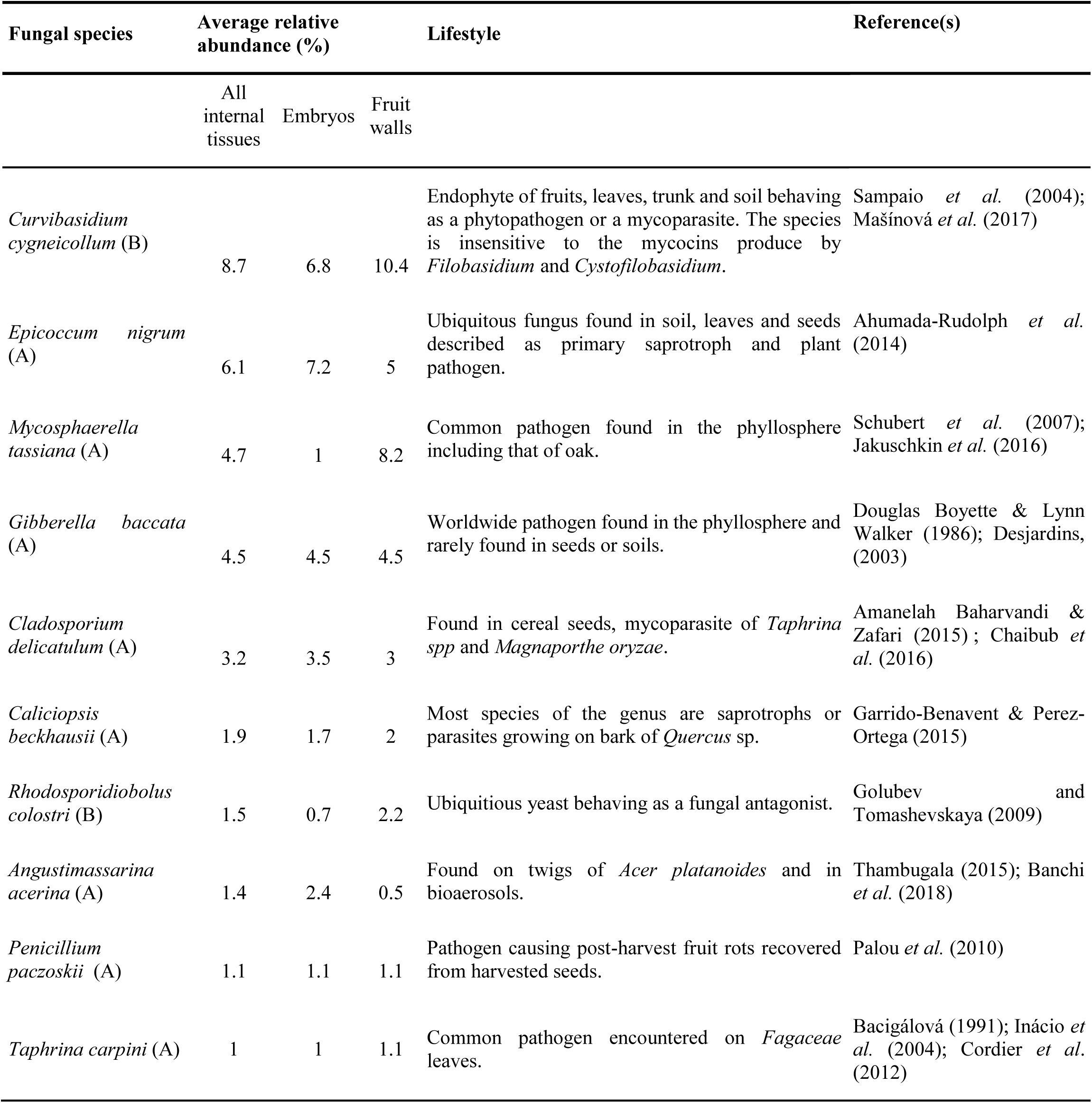
Most abundant fungal species found as endophytes in seeds of sessile oak. Seeds were collected in the tree canopy or on the ground and were surface sterilized. The fruit wall and embryo were separated and their fungal community was analyzed using a metabarcoding approach. Only Amplicon sequence variants (ASV) assigned to Ascomycota (A) or Basidiomycota (B) with the UNITE database were kept. Average relative abundances of all ASVs were computed for each sample type, after merging ASVs assigned to the same fungal species. Only ASVs identified at the species level are shown in the table.

| Fungal species | Average relative abundance (%) |  |  | Lifestyle | Reference(s) |
| --- | --- | --- | --- | --- | --- |
|  | All internal tissues | Embryos | Fruit walls |  |  |
| *Curvibasidium cygneicollum* (B) | 8.7 | 6.8 | 10.4 | Endophyte of fruits, leaves, trunk and soil behaving as a phytopathogen or a mycoparasite. The species is insensitive to the mycocins produce by *Filobasidium* and *Cystofilobasidium*. | Sampaio *et al.* (2004); Mašínová *et al.* (2017) |
| *Epicoccum nigrum* (A) | 6.1 | 7.2 | 5 | Ubiquitous fungus found in soil, leaves and seeds described as primary saprotroph and plant pathogen. | Ahumada-Rudolph *et al.* (2014) |
| *Mycosphaerella tassiana* (A) | 4.7 | 1 | 8.2 | Common pathogen found in the phyllosphere including that of oak. | Schubert *et al.* (2007); Jakuschkin *et al.* (2016) |
| *Gibberella baccata* (A) | 4.5 | 4.5 | 4.5 | Worldwide pathogen found in the phyllosphere and rarely found in seeds or soils. | Douglas Boyette & Lynn Walker (1986); Desjardins, (2003) |
| *Cladosporium delicatulum* (A) | 3.2 | 3.5 | 3 | Found in cereal seeds, mycoparasite of *Taphrina* spp and *Magnaporthe oryzae*. | Amanelah Baharvandi & Zafari (2015); Chaibub *et al.* (2016) |
| *Caliciopsis beckhausii* (A) | 1.9 | 1.7 | 2 | Most species of the genus are saprotrophs or parasites growing on bark of *Quercus* sp. | Garrido-Benavent & Perez-Ortega (2015) |
| *Rhodosporidiobolus colostri* (B) | 1.5 | 0.7 | 2.2 | Ubiquitous yeast behaving as a fungal antagonist. | Golubev and Tomashevskaya (2009) |
| *Angustimassarina acerina* (A) | 1.4 | 2.4 | 0.5 | Found on twigs of *Acer platanoides* and in bioaerosols. | Thambugala (2015); Banchi *et al.* (2018) |
| *Penicillium paczoskii* (A) | 1.1 | 1.1 | 1.1 | Pathogen causing post-harvest fruit rots recovered from harvested seeds. | Palou *et al.* (2010) |
| *Taphrina carpini* (A) | 1 | 1 | 1.1 | Common pathogen encountered on *Fagaceae* leaves. | Bacigálová (1991); Inácio *et al.* (2004); Cordier *et al.* (2012) |

**Table S2.** Permutational multivariate analyses of variance (PERMANOVA) of compositional dissimilarities among fungal communities of seeds of sessile oak. Dissimilarities among seeds were estimated using the Jaccard quantitative distance. The total number of sequences per sample (sequencing depth, SD) was log-transformed and introduced as the first explanatory variable in all models. The effects of tree population (T), seed position (canopy *versus* ground, P) and their interaction were tested on the whole acorn dataset while the effect of mother tree was tested separately on canopy seeds and ground seeds.

|  | d.f. | F-value | P-value | R<sup>2</sup> |
| --- | --- | --- | --- | --- |
| **All seeds** |  |  |  |  |
| Log(SD) | 1 | 1.3 | 0.018 | 0.01630 |
| Tree population (T) | 3 | 2.1 | **<.001** | 0.07967 |
| Seed position (P) | 1 | 1.8 | **<.001** | 0.02240 |
| T x P | 3 | 1.2 | **<.01** | 0.04619 |
| **Seeds in the canopy** |  |  |  |  |
| Log(SD) | 1 | 1.0 | 0.626 | 0.02588 |
| Mother tree | 10 | 1.2 | **0.011** | 0.30870 |
| **Seeds on the ground** |  |  |  |  |
| Log(SD) | 1 | 1.1 | 0.238 | 0.02935 |
| Mother tree | 11 | 1.2 | 0.150 | 0.34904 |

**Table S3.** Response of fungal ASVs to site elevation and seed position depending on their lifestyle, in the HMSC presence-absence (PA) and sequence count (SC) models. In the PA model, values are the posterior probabilities that the community weighted mean specialization, or the proportion of pathogens or saprotrophs is higher in the reference modality of the ecological factor. In the SC model, values are the posterior probabilities that the community weighted mean specialization, or the abundance of pathogens or saprotrophs (conditional on presence) is higher in the reference modality of the ecological factor. Probabilities higher than 0.9 are in bold.

| Ecological factor | Reference modality | PA model |  |  | SC model |  |  |
| --- | --- | --- | --- | --- | --- | --- | --- |
|  |  | Specialization | Pathogen | Saprotroph | Specialization | Pathogen | Saprotroph |
| *Elevation* | High | **0.96875** | 0.57875 | **0.97825** | 0.71475 | 0.84475 | 0.75225 |
| *Seed position* | Ground | 0.78575 | **0.92525** | **0.9225** | 0.38525 | 0.42925 | 0.33575 |

**Table S4.**
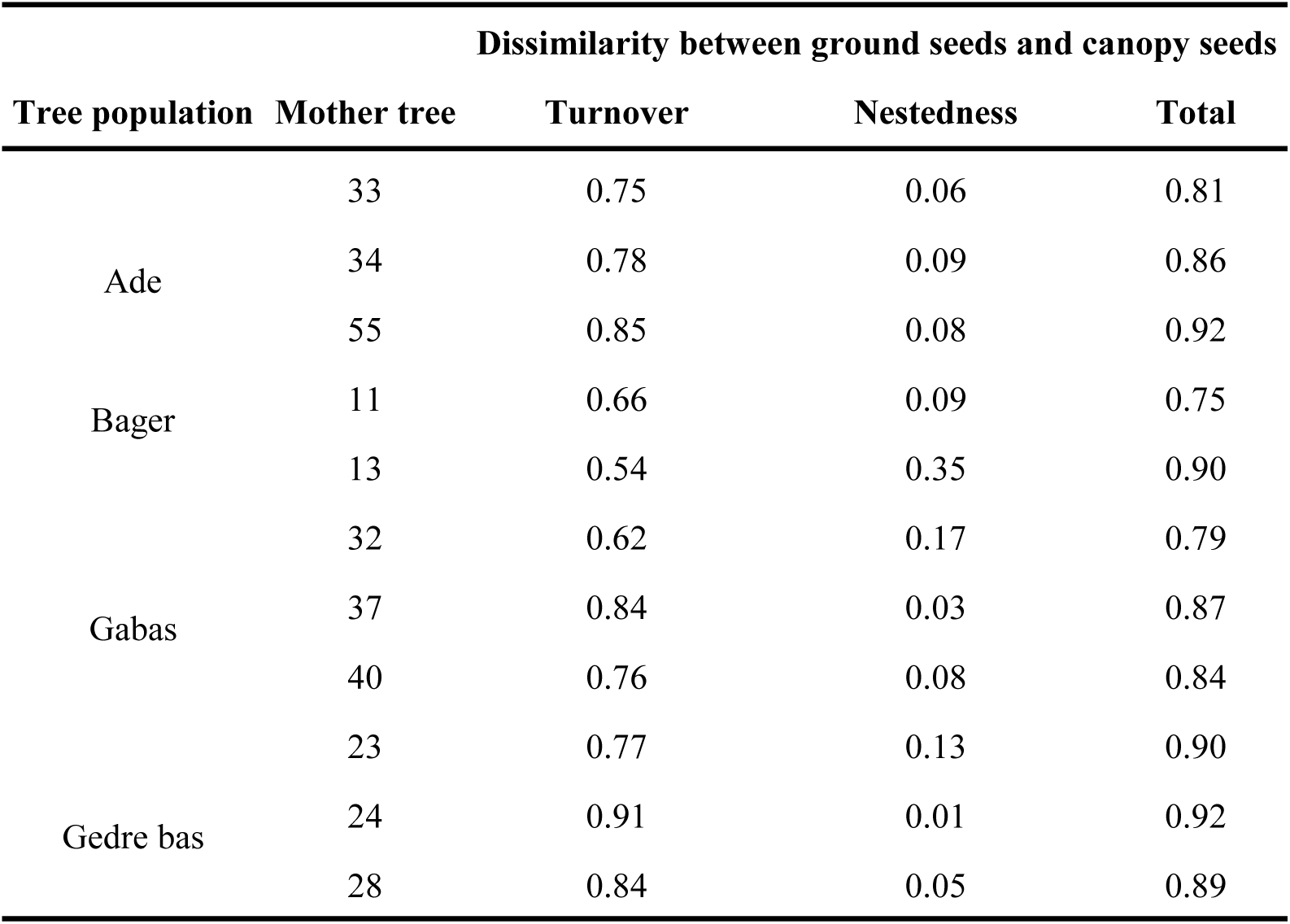
Nestedness and turnover components of compositional dissimilarities between fungal communities of seeds in the canopy and seeds on the ground. Compositional dissimilarities among fungal communities were estimated for each mother tree using the binary Jaccard dissimilarity index and partitioned with the beta.temp function of the betapart package (Baselga et al. 2018). A mother tree at the Bager site was excluded from the analysis because it had only two seeds on the ground.

**Table S5.**
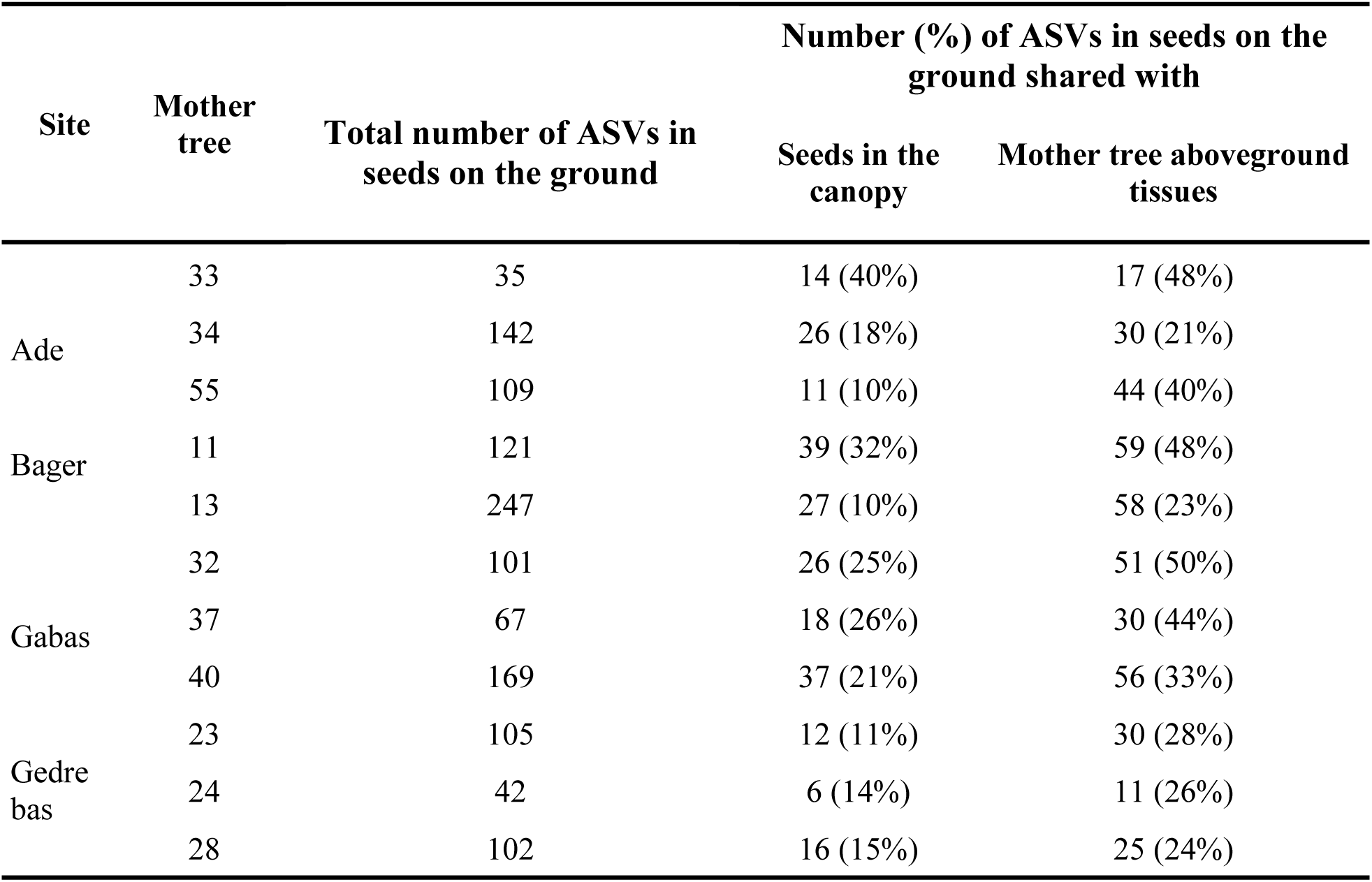
Number and percentage of fungal ASVs of seeds on the ground also found in seeds in the canopy and mother tree aboveground tissues. A mother tree at the Bager site was excluded from the analysis because it had only two acorns on the ground.

**Table S6.**
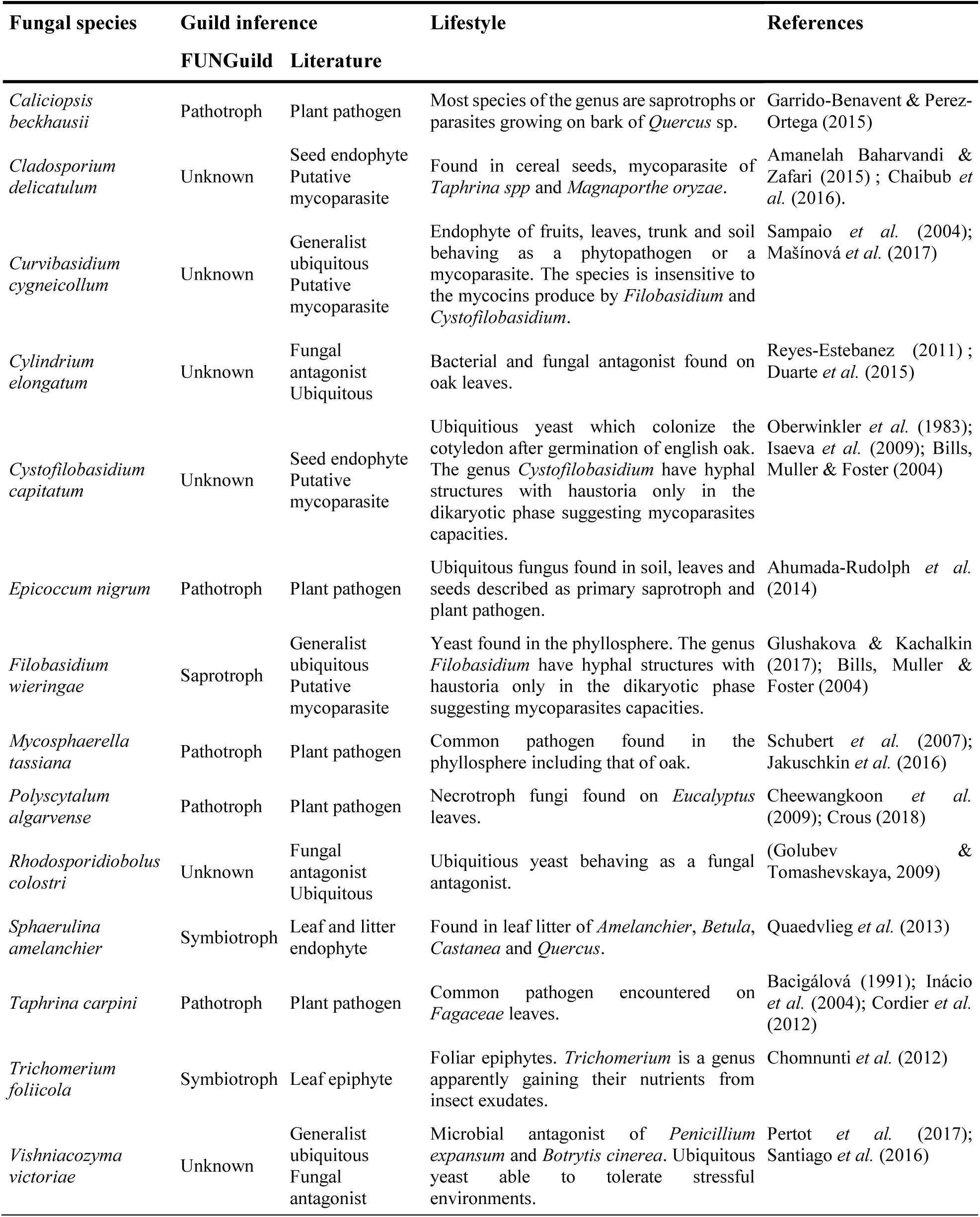
Fungal ASVs positively associated with another other ASV at the seed level according to HMSC models. Putative functions were inferred using FUNGuild and literature search. Only ASVs assigned at the species level are shown. The network of positive associations is represented on Figure 4.

**Figure S1.**
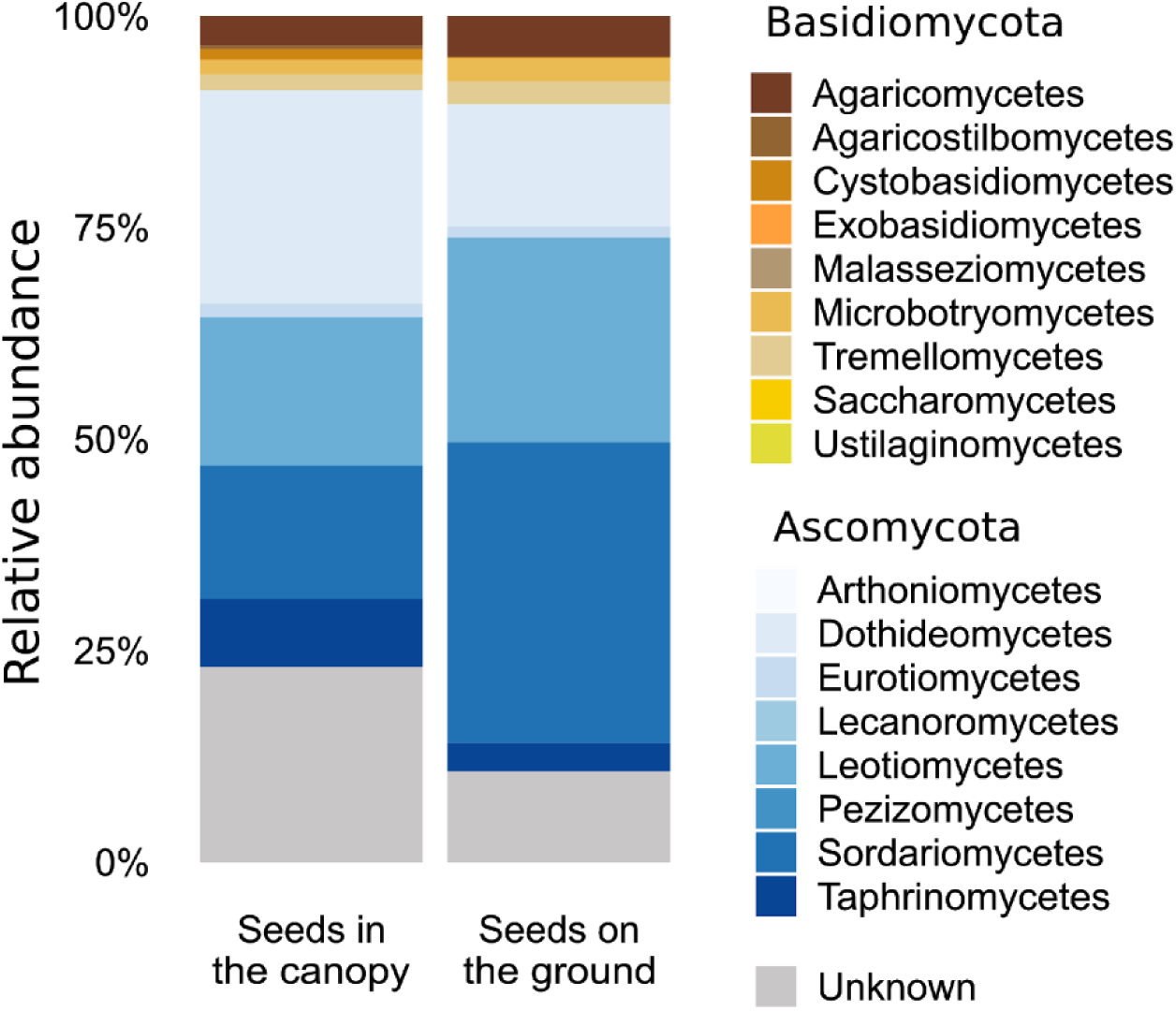
Taxonomic composition of fungal communities of seeds of sessile oak. The barplot indicates the average sequence percentage assigned to each fungal class.

**Figure S2.**
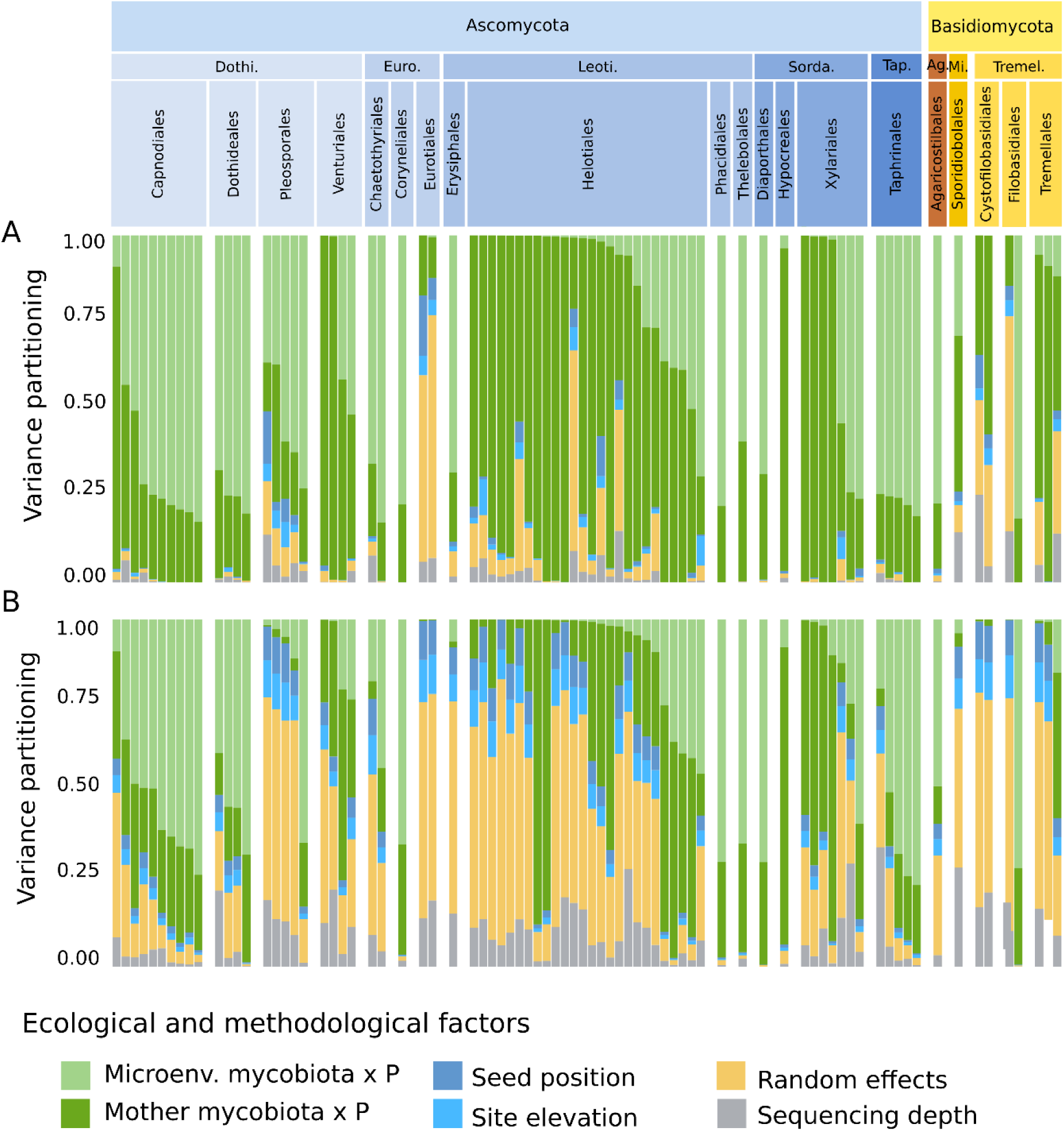
Partitioning of the variance in the composition of seed fungal communities of sessile oak. Four fixed effects were included in the HMSC models to explain variations in (A) the presence-absence or (B) the sequence count of a focal fungal ASV among seed samples: *Sequencing depth* (total number of sequences per sample), *Seed position* (P, canopy or ground), *Microenv. mycobiota* (relative abundance of the focal ASV in the seed microenvironment), *Mother mycobiota* (relative abundance of the focal ASV in the mother tree aboveground tissues), and *Site elevation. Mother mycobiota* and *Microenv. mycobiota* were introduced in interaction with P. Random effects were included at each spatial scale (seed, tree and population). Results of variance partitioning are given as percentages (%) of total explained variance. ASV are ranked by fungal phylum, class and order (Dothi: *Dothideomycetes*, Euro: *Eurotiomyecetes*, Leo: *Leotiomyecetes*, Sorda: *Sordariomycetes*, Tap: *Taphrinomycetes*, Ag: *Agaricostilbmoyecetes*, Mi: *Microbotryomycetes*, Tremel: *Tremellomycetes*). Only ASVs assigned at the order level are shown.

## References

Ahumada-Rudolph R, Cajas-Madriaga D, Rudolph A, Reinoso R, Torres C, Silva M, Becerra J. 2014. Variation of sterols and fatty acids as an adaptive response to changes in temperature, salinity and pH of a marine fungus *Epicoccum nigrum* isolated from the Patagonian Fjords. Revista de biología marina y oceanografía 49: 293–305.

Amanelah Baharvandi H, Zafari D. 2015. Identification of *Cladosporium delicatulum* as a mycoparasite of *Taphrina pruni*. Archives of Phytopathology and Plant Protection 48: 688–697.

Andrews JH, Harris RF. 2000. The ecology and biogeography of microorganisms on plant surfaces. Annual Review of Phytopathology 38: 145–180.

Bacigálová K. 1991. New localities of *Taphrina carpini* (Rostr.) Johans, on *Carpinus betulus* in Slovakia. Czech Mycology 46: 296–302.

Baker KF, Smith SH. 1966. Dynamics of seed transmission of plant pathogens. Annual Review of Phytopathology 4: 311 –332.

Barret M, Guimbaud JF, Darrasse A, Jacques MA. 2016. Plant microbiota affects seed transmission of phytopathogenic microorganisms. Molecular plant pathology 17: 791–795.

Baselga A, Orme CDL. 2012. Betapart: An R package for the study of beta diversity. Methods in Ecology and Evolution 3: 808–812.

Brader G, Compant S, Vescio K, Mitter B, Trognitz F, Ma L-J, Sessitsch A. 2017. Ecology and genomic insights into plant-pathogenic and plant-nonpathogenic endophytes. Annual Review of Phytopathology 55: 61–83.

Caignard T, Kremer A, Firmat C, Nicolas M, Venner S, Delzon S. 2017. Increasing spring temperatures favor oak seed production in temperate areas. Scientific Reports 7: 1–8.

Callahan BJ, McMurdie PJ, Rosen MJ, Han AW, Johnson AJA, Holmes SP. 2016. DADA2: High-resolution sample inference from Illumina amplicon data. Nature Methods 13: 581–583.

Campisano A, Ometto L, Compant S, Pancher M, Antonielli L, Yousaf S, Varotto C, Anfora G, Pertot I, Sessitsch A, et al. 2014. Interkingdom transfer of the acne-causing agent, propionibacterium acnes, from human to grapevine. Molecular Biology and Evolution 31: 1059–1065.

Cavazos BR, Bohner TF, Donald ML, Sneck ME, Shadow A, Omacini M, Rudgers JA, Miller TEX. 2018. Testing the roles of vertical transmission and drought stress in the prevalence of heritable fungal endophytes in annual grass populations. New Phytologist: 1075–1084.

Chaibub AA, de Carvalho JCB, de Sousa Silva C, Collevatti RG, Gonçalves FJ, de Carvalho Barros Côrtes MV, de Filippi MCC, de Faria FP, Lopes DCB, de Araújo LG. 2016. Defence responses in rice plants in prior and simultaneous applications of *Cladosporium* sp. during leaf blast suppression. Environmental Science and Pollution Research 23: 21554–21564.

Chancerel E, Lamy JB, Lesur I, Noirot C, Klopp C, Ehrenmann F, Boury C, Provost G Le, Label P, Lalanne C, et al. 2013. High-density linkage mapping in a pine tree reveals a genomic region associated with inbreeding depression and provides clues to the extent and distribution of meiotic recombination. BMC Biology 11: 50.

Chase JM, Myers JA. 2011. Disentangling the importance of ecological niches from stochastic processes across scales. Philosophical Transactions of the Royal Society 366: 2351–2363.

Cheewangkoon R, Groenewald JZ, Summerell BA, Hyde KD, To-Anun C, Crous PW. 2009. Myrtaceae, a cache of fungal biodiversity. Persoonia: Molecular Phylogeny and Evolution of Fungi 23: 55–85.

Chen H, Wu H, Yan B, Zhao H, Liu F, Zhang H, Sheng Q, Miao F, Liang Z. 2018. Core microbiome of medicinal plant *Salvia miltiorrhiza* seed: A rich reservoir of beneficial microbes for secondary metabolism? Molecular Sciences 19: 672–687.

Chomnunti P, Bhat DJ, Jones EBG. 2012. Trichomeriaceae, a new sooty mould family of Chaetothyriales. Fungal Diversity 56:63–76

Clay K, Schardl C. 2002. Evolutionary origins and ecological consequences of endophyte symbiosis with grasses. The American Naturalist 160: S99–S127.

Compant S, Van Der Heijden MGA, Sessitsch A. 2010. Climate change effects on beneficial plant-microorganism interactions. FEMS Microbiology Ecology 73: 197–214.

Coince A, Cordier T, Lengellé J, Defossez E, Vacher C, Robin C, Buée M, Marçais B. 2014. Leaf and root-associated fungal assemblages do not follow similar elevational diversity patterns. PLoS ONE 9.

Cordier T, Robin C, Capdevielle X, Fabreguettes O, Desprez-Loustau ML, Vacher C. 2012. The composition of phyllosphere fungal assemblages of European beech (*Fagus sylvatica*) varies significantly along an elevation gradient. New Phytologist 196: 510–519.

Crist T, Friese F. 2009. The Impact of Fungi on Soil Seeds: Implications for plants and granivores in a semiarid shrub-steppe. Ecological Society of America 74: 2231–2239.

Crous PW, Schumacher RK, Wingfield MJ, Akulov A, Denman S, Roux J, Braun U, Burgess TI, Carnegie AJ, Váczy KZ, et al. 2018. New and Interesting Fungi. Fungal Systematics and Evolution 1: 169–215.

Desjardins AE. 2003. Gibberella from A(venaceae) to Z(eae). Annual Review of Phytopathology 41: 177–198.

Douglas Boyette C, Lynn Walker H. 1986. Evaluation of *Fusarium lateritium* as a Biological Herbicide for Controlling Velvetleaf (Abutilon theophrasti) and Prickly Sida (Sida spinosa). Weed Science Society of America 34: 106–109.

Duarte S, Bärlocher F, Trabulo J, Cássio F. 2015. Stream-dwelling fungal decomposer communities along a gradient of eutrophication unraveled by 454 pyrosequencing. Fungal Diversity 70: 127–148.

Escobar Rodriguez C, Mitter B, Barret M, Sessitsch A, Compant S. 2018. Commentary: seed bacterial inhabitants and their routes of colonization. Plant and Soil 422: 129–134.

Ewald PW. 1989. Transmission modes and evolution of the parasitism-mutualism continuum. Annals of the New York Academy of Sciences 503: 295–306.

Faeth SH. 2002. Fungal Endophytes: Common Host Plant Symbionts but Uncommon Mutualists. Integrative and Comparative Biology 42: 360–368.

Feldman TS, Brien HEO, Arnold AE. 2008. Moths that vector a plant pathogen also transport endophytic fungi and mycoparasitic antagonists. Microbial Ecology 56: 742–750.

Frank A, Saldierna Guzmán J, Shay J. 2017. Transmission of bacterial endophytes. Microorganisms 5: 70.

Galan M, Razzauti M, Bard E, Bernard M, Brouat C, Charbonnel N, Dehne-Garcia A, Loiseau A, Tatard C, Tamisier L, et al. 2016. 16S rRNA amplicon sequencing for epidemiological surveys of bacteria in wildlife. mSystems 1: e00032–16.

Gardes M, Bruns TD. 1993. ITS primers with enhanced specificity for basidiomycetes - application to the identification of mycorrhizae and rusts. Molecular Ecology 2: 113–118.

Garrido-benavent I, Pérez-ortega S. 2015. Unravelling the diversity of European Caliciopsis (Coryneliaceae, Ascomycota): *Caliciopsis valentina* sp. nov. and *C. beckhausii* comb. nov., with a worldwide key to Caliciopsis. Mycological Progress 14: 1–11.

Gerzabek G, Oddou-Muratorio S, Hampe A. 2017. Temporal change and determinants of maternal reproductive success in an expanding oak forest stand. Journal of Ecology 105: 39–48.

Gilbert GS. 2002. Evolutionary ecology of plant diseases in natural ecosystems. Annual Review of Phytopathology 40: 13–43.

Glassner H, Zchori-Fein E, Compant S, Sessitsch A, Katzir N, Portnoy V, Yaron S. 2015. Characterization of endophytic bacteria from cucurbit fruits with potential benefits to agriculture in melons (Cucumis melo L.). FEMS Microbiology Ecology 91: 1–13.

Glushakova AM, Kachalkin A V. 2017. Endophytic yeasts in *Malus domestica* and *Pyrus communis* fruits under anthropogenic impact. Microbiology 86: 128–135.

Golubev WI, Tomashevskaya MA. 2009. Characterization of Mycocin Secreted by *Rhodotorula colostri* (Castelli) Lodder. Biology bulletin 36: 373–378.

Gundel PE, Rudgers JA, Whitney KD. 2017. Vertically transmitted symbionts as mechanisms of transgenerational effects. American Journal of Botany 104: 787–792.

Jaccard P. 1901. Étude comparative de la distribution florale dans une portion des Alpes et du Jura. Bulletin de la Société Vaudoise des Sciences Naturelles 37: 547–579.

Inácio J, Rodrigues MG, Sobral P, Fonseca Á. 2004. Characterisation and classification of phylloplane yeasts from Portugal related to the genus *Taphrina* and description of five novel *Lalaria* species. FEMS Yeast Research 4: 541–555.

Isaeva O V., Glushakova AM, Yurkov AM, Chernov IY. 2009. The yeast *Candida railenensis* in the fruits of English oak (Quercus robur L.). Microbiology 78: 355–359.

Jakuschkin B, Fievet V, Schwaller L, Fort T, Robin C, Vacher C. 2016. Deciphering the pathobiome: Intra- and interkingdom interactions involving the pathogen *Erysiphe alphitoides*. Microbial Ecology 72: 870–880.

Janzen DH. 1970. Herbivores and the Number of Tree Species in Tropical Forests. The American Naturalist 104: 501–528.

Johnston-Monje D, Raizada MN. 2011. Conservation and diversity of seed associated endophytes in Zea across boundaries of evolution, ethnography and ecology. PLoS ONE 6:e20396.

Klaedtke S, Jacques MA, Raggi L, Préveaux A, Bonneau S, Negri V, Chable V, Barret M. 2016. Terroir is a key driver of seed-associated microbial assemblages. Environmental microbiology 18: 1792–1804.

Leff JW, Lynch RC, Kane NC, Fierer N. 2017. Plant domestication and the assembly of bacterial and fungal communities associated with strains of the common sunflower, *Helianthus annuus*. New Phytologist 214: 412–423.

Leroy C, Maes AQ, Louisanna E, Séjalon-Delmas N. 2019. How significant are endophytic fungi in bromeliad seeds and seedlings ? Effects on germination, survival and performance of two epiphytic plant species. Fungal Ecology 39: 296–306.

Love MI, Huber W, Anders S, Love MI, Huber W, Anders S. 2014. Moderated estimation of fold change and dispersion for RNA-Seq data with DESeq2. Genome biology 15: 521–550.

Mašínová T, Bahnmann BD, Větrovský T, Tomšovský M, Merunková K, Baldrian P. 2017. Drivers of yeast community composition in the litter and soil of a temperate forest. FEMS microbiology ecology 93: 1–10.

Maude RB. 1996. Seedborne Diseases and Their Control: Principles and Practice.

McMurdie PJ, Holmes S. 2013. Phyloseq: An R Package for Reproducible Interactive Analysis and Graphics of Microbiome Census Data. PLoS ONE 8: e61217.

Mcshea WJ, Healy WM, Devers P, Fearer T, Frank H, Stauffer D, Waldon J. 2019. Forestry matters: Decline of oaks will impact wildlife in hardwood forests. Journal of Wildlife management 71: 1717–1728.

Minard G, Tikhonov G, Ovaskainen O, Saastamoinen M. Variation in Melitaea cinxia gut microbiota is phylogenetically highly structured but 2only mildly driven by host plant microbiota, sex or parasitism. bioRxiv, in press.

Monahan WB, Koenig WD. 2005. Estimating the potential effects of sudden oak death on oak-dependent birds. Biological Conservation 127: 146–157.

Mueller, G.M., Bills, G.F., Foster MSB. 2004. Biodiversity of Fungi: Inventory and monitoring methods.

Nalim FA, Samuels GJ, Wijesundera RL, Geiser DM. 2011. New species from the *Fusarium solani* species complex derived from perithecia and soil in the old world tropics. Mycologia 103: 1302–1330.

Nelson EB. 2004. Microbial dynamics and interactions in the spermosphere. Annual Review of Phytopathology 42: 271–309.

Nelson EB. 2017. The seed microbiome: origins, interactions, and impacts. Plant and Soil 422: 7–34.

Nelson EB, Simoneau P, Barret M, Mitter B, Compant S. 2018. The soil, the seed, the microbes and the plant. Plant Soil 422: 1–5.

Nemergut DR, Schmidt SK, Fukami T, O’Neill SP, Bilinski TM, Stanish LF, Knelman JE, Darcy JL, Lynch RC, Wickey P, et al. 2013. Patterns and processes of microbial community assembly. Microbiology and Molecular Biology Reviews 77: 342–356.

Nguyen NH, Song Z, Bates ST, Branco S, Tedersoo L, Menke J, Schilling JS, Kennedy PG. 2016. FUNGuild: an open annotation tool for parsing fungal community datasets by ecological guild. Fungal Ecology 20: 241–248.

Oberwinkler F, Bandoni R, Blanz P. 1983. Cystofilobasidium: a New Genus in the *Filobasidiaceae* *. Systematic and applied microbiology 4: 114–122.

Ofek M, Hadar Y, Minz D. 2011. Colonization of cucumber seeds by bacteria during germination. Environmental Microbiology 13: 2794–2807.

Ovaskainen O, Tikhonov G, Norberg A, Guillaume Blanchet F, Duan L, Dunson D, Roslin T, Abrego N. 2017. How to make more out of community data? A conceptual framework and its implementation as models and software. Ecology Letters 20: 561–576.

Palou L, Guardado A, Monstesinos-Herrero C. 2010. First report of *Penicillium* spp. and *Pilidiella granati* causing postharvest fruit rot of pomegranate in Spain. New disease reports 22: 21.

Peris JE, Rodríguez A, Peña L, Fedriani JM. 2017. Fungal infestation boosts fruit aroma and fruit removal by mammals and birds. Scientific Reports: 1–9.

Pertot I, Giovannini O, Benanchi M, Caf T, Rossi V, Mugnai L. 2017. Combining biocontrol agents with different mechanisms of action in a strategy to control Botrytis cinerea on grapevine. 97: 85–93.

Pinciroli M, Gribaldo A, Vidal A, Bezus R, Sisterna M. 2013. Mycobiota evolution during storage of paddy, brown and milled rice in different genotypes. Summa phytopathologica 39: 157–161.

Prochazkova Z, Sikorova A, Peskova V. 2005. Preliminary observations on the occurrence of *Ciboria batschiana* (Zopf) Buchwald in the Czech Republic. Working Papers of the Finnish Forest Research Institute 11: 13–18.

Qin Y, Pan X, Yuan Z. 2016. Seed endophytic microbiota in a coastal plant and phytobeneficial properties of the fungus *Cladosporium cladosporioides*. Fungal Ecology 24: 53–60.

Quaedvlieg W, Verkley GJM, Shin HD, Barreto RW, Alfenas AC, Swart WJ, Groenewald JZ, Crous PW. 2013. Sizing up *Septoria*. Studies in Mycology 75: 307–390.

Reyes-Estebanez M. 2011. Antimicrobial and nematicidal screening of anamorphic fungi isolated from plant debris of tropical areas in Mexico. African Journal of Microbiology Research 5: 1083–1089.

Rezki S, Campion C, Simoneau P, Jacques MA, Shade A, Barret M. 2018. Assembly of seed-associated microbial communities within and across successive plant generations. Plant and Soil 422: 67–79.

Rillig MC, Antonovics J, Caruso T, Lehmann A, Powell JR, Veresoglou SD, Verbruggen E. 2015. Interchange of entire communities: Microbial community coalescence. Trends in Ecology and Evolution 30: 470–476.

Sampaio JP, Golubev WI, Fell JW, Gadanho M, Golubev NW. 2004. *Curvibasidium cygneicollum* gen. nov., sp. nov. and *Curvibasidium pallidicorallinum* sp. nov., novel taxa in the microbotryomycetidae (Urediniomycetes), and their relationship with *Rhodotorula fujisanensis* and *Rhodotorula nothofagi*. International Journal of Systematic and Evolutionary Microbiology 54: 1401–1407.

Santiago IF, Rosa CA, Rosa LH. 2016. Endophytic symbiont yeasts associated with the Antarctic angiosperms Deschampsia antarctica and Colobanthus quitensis. Polar Biology 40:177–183

Schiltz S, Gaillard I, Pawlicki-Jullian N, Thiombiano B, Mesnard F, Gontier E. 2015. A review: what is the spermosphere and how can it be studied? Journal of Applied Microbiology 119: 1467–1481.

Schneider C, Rasband W, Eliceiri K. 2012. NIH Image to ImageJ: 25 years of image analysis. Nature Methods 9: 671–675.

Schoch CL, Seifert KA, Huhndorf S, Robert V, Spouge JL, Levesque CA, Chen W, Bolchacova E, Voigt K, Crous PW, et al. 2012. Nuclear ribosomal internal transcribed spacer (ITS) region as a universal DNA barcode marker for Fungi. Proceedings of the National Academy of Sciences 109: 6241–6246.

Schubert K, Groenewald JZ, Braun U, Dijksterhuis J, Starink M, Hill CF, Zalar P, De Hoog GS, Crous PW. 2007. Biodiversity in the *Cladosporium herbarum* complex (*Davidiellaceae, Capnodiales*), with standardisation of methods for *Cladosporium* taxonomy and diagnostics. Studies in Mycology 58: 105–156.

Shahzad R, Khan AL, Bilal S, Asaf S, Lee I. 2018. What Is There in Seeds? Vertically Transmitted Endophytic Resources for Sustainable Improvement in Plant Growth. Frontiers in Plant Science 9.

Sneck ME, Rudgers JA, Young CA, Miller TEX. 2017. Variation in the prevalence and Transmission of heritable symbionts across host populations in heterogeneous environments. Microbial Ecology 74: 640–653.

Sogonov M V., Castlebury LA, Rossman AY, Mejía LC, White JF. 2008. Leaf-inhabiting genera of the *Gnomoniaceae, Diaporthales*. Studies in Mycology 62: 1–77.

Thambugala KM, Hyde KD, Tanaka K, Tian Q, Wanasinghe DN, Ariyawansa HA, Jayasiri SC, Boonmee S, Camporesi E, Hashimoto A, et al. 2015. Towards a natural classification and backbone tree for Lophiostomataceae, Floricolaceae, and Amorosiaceae fam. nov. Fungal Diversity 74: 199–266.

Torres-Cortés G, Bonneau S, Bouchez O, Genthon C, Briand M, Jacques M-A, Barret M. 2018. Functional microbial features driving community assembly during seed germination and emergence. Frontiers in Plant Science 9: 902.

Truyens S, Weyens N, Cuypers A, Vangronsveld J. 2015. Bacterial seed endophytes: Genera, vertical transmission and interaction with plants. Environmental Microbiology Reports 7: 40–50.

U’Ren JM, Arnold AE. 2016. Diversity, taxonomic composition, and functional aspects of fungal communities in living, senesced, and fallen leaves at five sites across North America. PeerJ 4: e2768.

Vacher C, Hampe A, Porté AJ, Sauer U, Compant S, Morris CE. 2016. The Phyllosphere: Microbial jungle at the plant–climate interface. Annual Review of Ecology, Evolution, and Systematics 47: 1–24.

Vannier N, Mony C, Bittebiere A, Michon-coudouel S, Biget M, Vandenkoornhuyse P. 2018. A microorganisms’ journey between plant generations. Microbiome 6: 1–11.

Vellend M. 2010. Conceptual synthesis in community ecology. The quarterly review of biology 85: 183–206.

Vivas M, Kemler M, Slippers B. 2015. Maternal effects on tree phenotypes: Considering the microbiome. Trends in Plant Science 20: 541–544.

Vivas M, Kemler M, Mphahlele MM, Wingfield MJ, Slippers B. 2017. Maternal effects on phenotype, resistance and the structuring of fungal communities in *Eucalyptus grandis*. Environmental and Experimental Botany 140: 120–127.

Wang Q, Garrity GM, Tiedje JM, Cole JR. 2007. Naïve Bayesian classifier for rapid assignment of rRNA sequences into the new bacterial taxonomy. Applied and Environmental Microbiology 73: 5261–5267.

Whitham TG, Bailey JK, Schweitzer| JA, Shuster SM, Bangert RK, LeRoy CJ, Lonsdorf E V., Allan GJ, DiFazio SP, Potts BM, et al. 2006. A framework for community and ecosystem genetics: from genes to ecosystems. Nature Reviews Genetics 7: 510–523.

Wolf JB, Wade MJ. 2009. What are maternal effects (and what are they not)? 364: 1107–1115.

Yang L, Danzberger J, Schöler A, Schröder P, Schloter M, Radl V. 2017. Dominant Groups of Potentially Active Bacteria Shared by Barley Seeds become Less Abundant in Root Associated Microbiome. Frontiers in Plant Science 8: 1–12.

Zhang J, Kobert K, Flouri T, Stamatakis A. 2014. PEAR: A fast and accurate Illumina Paired-End reAd mergeR. Bioinformatics 30: 614–620.

Zhou J, Ning D. 2017. Stochastic community assembly: Does it matter in microbial ecology? Microbiology and Molecular Biology Reviews 81: 2–17.

